# Molecular evolution of the Wood-Ljungdahl pathway and the reductive glycine pathway in Thermodesulfobacteriota

**DOI:** 10.1101/2025.09.01.672775

**Authors:** Tomoyuki Wakashima, Keitaro Kume, Yoko Chiba

## Abstract

Carbon fixation is a fundamental metabolic process that sustains ecosystems, yet its origins and evolutionary history remain largely unresolved. In this study, we focused on the Wood-Ljungdahl (WL) pathway, which is considered one of the most ancient carbon fixation pathways and the reductive glycine (rGly) pathway, which shares several reactions with the WL pathway. The evolutionary scenario of the two carbon fixation pathways was inferred in the phylum Thermodesulfobacteriota, which includes microorganisms that operate either the WL pathway or the rGly pathway for autotrophic growth. The timing of gene gain and loss events was inferred by gene presence/absence analyses for both pathways, together with phylogenetic analyses of their key enzymes. Our results suggested that the common ancestor of Thermodesulfobacteriota possessed all genes encoding key enzymes of both pathways; formate dehydrogenase, the carbon monoxide dehydrogenase/acetyl-CoA synthase complex and the glycine cleavage system. Furthermore, analyses of complete gene sets for the WL and rGly pathway, together with downstream genes required for amino acid biosynthesis, supported the possibility that the common ancestor of this phylum had been capable of autotrophic growth through these carbon fixation pathways. Then, multiple lineages have lost the WL and rGly pathway genes independently during subsequent evolution. Gene replacements also occurred in the glycine cleavage system by regaining genes by horizontal gene transfer. These results suggest that carbon fixation pathways in extant organisms in the phylum Thermodesulfobacteriota arose through a combination of vertical inheritance, gene loss, and horizontal gene transfer.

## 1 Introduction

Elucidating the origins and early evolution of carbon fixation pathways is essential to understand the origin of life. Whether the first organisms on Earth were autotrophs or heterotrophs remains unresolved (Bada and Lazcano, 2002; Kitadai and Maruyama, 2018; Wächtershäuser, 1988; Kitadai et al., 2021), however, there is no doubt that the emergence of autotrophs was crucial for establishing ecosystem on Earth. This is because organic compounds produced solely through abiotic processes are insufficient to sustain a substantial biomass (Martin and Russell, 2007; Nunoura et al., 2018; Weiss et al., 2016; Berg et al., 2010; Dick, 2019). There is growing interest in how early life fixed carbon dioxide (CO_2_) and how it evolved to acquire the complex and highly refined metabolic systems observed today (Fuchs, 2011; Braakman and Smith, 2012; Moody et al., 2024).

To date, seven carbon fixation pathways have been identified in extant organisms in nature (Berg et al., 2010; Fuchs, 2011; Sánchez-Andrea et al., 2020). Among these pathways, the Wood-Ljungdahl (WL) pathway, which is also called the reductive acetyl-CoA pathway, is considered to be one of the oldest ones and existed in the last universal common ancestor (LUCA), owing to its broad conservation across diverse lineages including bacteria and archaea (Martin and Russell, 2007; Weiss et al., 2016; Berg et al., 2010; Braakman and Smith, 2012; Schuchmann and Müller, 2014; Moody et al., 2024; Nitschke and Russell, 2013; Peretó et al., 1999). The old origin of the WL pathway is also supported by its structural simplicity and high energy efficiency (Martin and Russell, 2007; Berg et al., 2010; Fuchs, 2011; Schuchmann and Müller, 2014).

The WL pathway incorporates two molecules of CO_2_ through the methyl and the carbonyl branches to synthesize one molecule of acetyl-CoA (Fig. 1, highlighted in blue) (Schuchmann and Müller, 2014). In the methyl branch of the bacterial WL pathway, CO_2_ is converted to formate by formate dehydrogenase (FDH) and subsequently reduced to a methyl group (-CH_3_) bound to tetrahydrofolate (THF) to produce CH_3_-THF. In the carbonyl branch, CO_2_ is reduced to CO by carbon monoxide dehydrogenase (CODH) and then combines with the methyl group of the CH_3_-THF and CoA to become acetyl-CoA by acetyl-CoA synthase (ACS). CODH and ACS involved in the WL pathway form a complex composed of five subunits; AcsA, AcsB, AcsC, and AcsD, are conserved between bacteria and archaea and AcsE and CdhB are specific to bacteria and archaea, respectively (Supplemental Table. S1). In both bacteria and archaea, the genes for these five subunits form a cluster (Adam et al., 2018). Both genetic and biochemical studies have confirmed that this pathway actually functions for carbon fixation in a wide range of taxa, including methanogens, acetogens, and sulfate-reducing bacteria, spanning both archaeal and bacterial domains (Schuchmann and Müller, 2014; Adam et al., 2018; Borrel et al., 2016; Pierce et al., 2008).

**Fig. 1.**
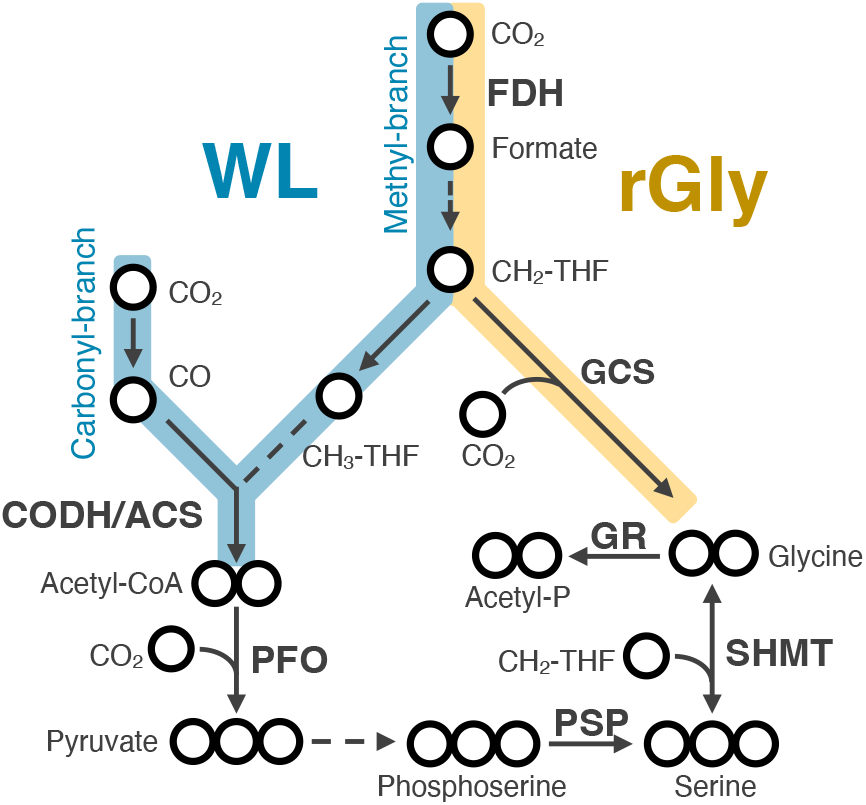
Metabolic map of the Wood–Ljungdahl (WL) and reductive glycine (rGly) pathways. Blue and yellow arrows indicate reactions in the WL and rGly pathways, respectively. Arrows represent enzymatic reactions and dashed arrows indicate reactions catalyzed by multiple enzymes. Bold letters show enzyme names. Circles represent number of carbons. THF: tetrahydrofolate; FDH: formate dehydrogenase; CODH/ACS: carbon monoxide dehydrogenase/acetyl-CoA synthase; GCS: glycine cleavage system; PFO: pyruvate-ferredoxin oxidoreductase; SHMT: serine hydroxymethyltransferase; PSP: phosphoserine phosphatase; GR: glycine reductase.

The reductive glycine (rGly) pathway (Fig. 1, highlighted in yellow) is the most recently identified CO_2_ fixation pathway that supports the autotrophic growth of naturally existing organisms (Sánchez-Andrea et al., 2020; Song et al., 2020). This pathway utilizes the reverse reaction of the glycine cleavage system (GCS), which cleave glycine to methylene tetrahydrofolate (CH_2_-THF), CO_2_, and ammonia (Fig. 1, highlighted in yellow) (Kikuchi et al., 2008; Kikuchi, 1973; Kim et al., 2017). GCS is composed of five proteins named GCST, GCSPα, GCSPβ, GCSL, and GCSH, and their genes are known to form a cluster (Okamura-Ikeda et al., 1993; Cao et al., 2018; Tezuka and Ohnishi, 2014; Lorio et al., 2010). GCSP catalyzes the decarboxylation of glycine and the product, aminomethyl group (-CH_2_-NH_2_), is transferred to a carrier protein, GCSH. The aminomethyl group reacts with tetrahydrofolate (THF) to produce CH_2_-THF and ammonia, which is catalyzed by GCST. The released GCSH is then re-oxidizes by GCSL, a FAD-dependent oxidoreductase (Kikuchi et al., 2008; Kikuchi, 1973). Because the GCS reaction is thermodynamically near equilibrium (Kikuchi et al., 2008), glycine synthesis through the reverse GCS reaction to supplement organic compound has been known for a long time in some heterotrophs (Schneeberger, 1999; Dürre and Andreesen, 1982; Waber and Wood, 1979; Fuchs, 1986). On the other hand, biochemically demonstrated CO_2_ fixation to support the autotrophic growth via the reverse GCS reaction (i.e., the rGly pathway) is limited to two species; *Desulfovibrio desulfuricans* belonging to the phylum Thermodesulfobacteriota (TDB) (Sánchez-Andrea et al., 2020) and *Clostridium drakei* belonging to the phylum Bacillota (Song et al., 2020). Notably, the pathway to provide CH_2_-THF from CO_2_ is shared between the rGly and WL pathways (Fig. 1, S1) (Sánchez-Andrea et al., 2020), suggesting the profound functional and evolutionary relationships between them (Song et al., 2020; Cotton et al., 2018). In fact, *Clostridium drakei* has been biochemically proven to operate both the rGly and WL pathways for CO_2_ fixation (Song et al., 2020). Furthermore, last bacterial common ancestor (LBCA) is suggested to have retained some part of the genes for these metabolic pathways. Ancestral metabolic reconstruction based on the gene phylogenies across major bacterial lineages identified methyl-branch enzyme genes shared between the WL and rGly pathways in the LBCA, although it did not support the presence of FDH, which catalyzes the first reaction in these pathways (Coleman et al., 2021).

FDH working for the WL or rGly pathways forms a hetromeric protein complex (Maia et al., 2015). For example, the FDH in *Moorella thermoacetica*, a model bacterium for the WL pathway, has an α_2_β_2_ structure (Yamamoto et al., 1983). In contrast, the FDH in *D. desulfuricans*, which operates the rGly pathway, consists of three subunits; α, β, and γ (Maia et al., 2016). The FDHα subunit in both species contains a metal that binds to CO_2_ and belong to the MopB superfamily, however, they are catabolized into distinct groups; the FDHα subunits of *M. thermoacetica* and *D. desulfuricans* are categorized into cytoplasmic FDH and FdhG, respectively, based on the distinct clade on their phylogenetic tree (Wells et al., 2023). In contrast, FDHβ exhibit no homology between in *M. thermoacetica* and in *D. desulfuricans*, and therefore expected to have different evolutionary origins. The phylogenetic diversity of FDH makes it difficult to estimate the existence/absent of the gene in ancestral organisms based on phylogenetic analysis, because the estimation is influenced by the query, taxon sampling, and root positions (Coleman et al., 2021; Powell and Battistuzzi, 2022; Superson et al., 2019).

Here, we estimated the evolutionary scenario of the genes related to CO_2_ fixation pathways in a thermophilic and sulfate-reducing bacterial phylum Thermodesulfobacteriota (TDB). The phylum TDB, formerly classified as Deltaproteobacteria, belongs to the Gracilicutes within the bacterial domain (Coleman et al., 2021) and is phylogenetically related to the phyla Myxococcota, Bdellovibrionota, and Proteobacteria (Fig. S2) (Waite et al., 2020). TDB includes *Desulfovibrio desulfuricans* and *Thermodesulfatator indicus* growing chemolithoautotrophically using the rGly and WL pathways, respectively (Sánchez-Andrea et al., 2020; Chiba et al., 2025). Therefore, this phylum is suitable to estimate the presence or absence of the gene set for the rGly and WL pathways in the common ancestor of this phylum and how it evolved to the extant organisms which have either pathway. To access these questions, we conducted presence/absence analyses for both pathway genes to identify the gene sets in extant organisms and performed phylogenetic analyses to estimate how the genes have been inherited.

## 2 Materials and Methods

### 2.1 Gene presence/absence analysis

To examine the gene presence/absence, we selected at least one representative species from each of the 42 families within to the phylum TDB based on the classification proposed by Waite et al. (Waite et al., 2020). Priority was given to those whose genomes are registered in RefSeq.

Proteomes for each representative species were predicted from NCBI RefSeq, and the presence or absence of enzymes involved in the WL and rGly, and subsequent amino acid synthesis pathways was estimated by homology searches using BLAST (Altschul et al., 1990). As queries, we used 39 amino acid sequences of the enzymes from *M. thermoacetica, D. desulfuricans, T. indicus, Escherichia coli, Thermus thermophilus* and *Hydrogenobacter thermophilus* (Supplementary Table 2). In the BLAST searches, we defined the homolog as those with an e-value of <1e-5 and query coverage of 70% or more.

In this study, we defined FDH, the CODH/ACS complex, and the GCS complex as the key enzymes of the WL and rGly pathways. We inferred that a species possessed the WL pathway if its genome encoded both a homolog of FDHα subunit and all five proteins of the CODH/ACS complex; AcsA, AcsB, AcsC, AcsD, and AcsE. We inferred that a species possessed the rGly pathway if its genome encoded homologs of FDHα and all the five GCS complex proteins; GCSPα, GCSPβ, GCST, GCSL, and GCSH.

### 2.2 Phylogenetic analyses

The target enzymes for phylogenetic analyses are as follows: FDH, the CODH/ACS complex, the GCS complex, pyruvate-ferredoxin oxidoreductase (PFO), phosphoserine phosphatase (PSP), serine hydroxymethyltransferase (SHMT), and the glycine reductase (GR) complex. Finally, a total of 28 enzyme subunits were the subjects of the analyses (Supplementary Table S3).

We collected sequences by MMseqs2 (release 14-7e284) similarity searches (Steinegger and Söding, 2017) against a local NCBI-nr database (ver. 2024/Nov/03). In these searches, we used sequences whose enzymatic functions were biochemically analyzed (Supplementary Table S3). The command line options of MMseqs2 search were “-e 1e-5 -s 7.0 --num-iterations 5”. From the result of the similarity search of each query, we first removed bottom 90% of MAG (metagenome-assembled genome)-derived hits by e-value. Then, we selected one representative sequence from each of NCBI taxid set we specified to cover the diversity of the tree of life. This set includes NCBI taxids of 199: 139 from whole organisms (for detail, see the legend to Supplementary Figure 1 in the previous study (Koganemaru et al., submitted)) and 60 related to TDB. These representatives on each query were finally merged. Next, the domain structures of the merged sequences were identified using InterProScan ver. 5.72, and sequences whose domain structures did not match those of the queries were removed (Jones et al., 2014). Sequences with large insertions or deletion were also removed to ensure accurate alignments. Next, in the preliminary phylogeny, we manually excluded sequences that exhibited long branches (>1.5 substitution per site) in the preliminary phylogenetic trees to minimize the effects of long-branch attraction (LBA) artefact (Bergsten, 2005; Graybeal, 1998; Hedtke et al., 2006). Because more stringent filtering criteria were applied in this phylogenetic analysis compared to the preceding gene presence/absence analysis to avoid the LBA artefact, some sequences that had been identified as “present” in the preceding analysis were excluded from the phylogenetic analysis. The numbers of sequences ultimately used and excluded are listed in Supplementary Table S3.

The curated sequences were aligned using MUSCLE (Edgar, 2004) and subsequently trimmed with G-blocks (Talavera and Castresana, 2007). The following three options were enabled for trimming: I. Allow smaller final blocks, II. Allow gap positions within the final blocks, and III. Allow less strict flanking positions. Phylogenetic trees were then inferred by maximum likelihood method using IQ-TREE ver. 2.3.6 on the final trimmed alignment data (Minh et al., 2020). Model selection was performed using ModelFinder “-m MFP” (Kalyaanamoorthy et al., 2017), which automatically determined the best-fit model for each enzyme. Node support values were evaluated using ultrafast bootstrap analysis “-B 1000” (Hoang et al., 2018). The evolutionary models and the number of selected amino acid sites used to infer each phylogenetic tree were provided in the legends of the respective figures. The workflow for the phylogenetic analysis is illustrated in Supplementary Figure S10.

For FDHα, a single phylogenetic tree was inferred by combining the newly collected homologous sequences of W-FDH (cytoplasmic Fdhs) from *M. thermoacetica* and Mo-FDH (FdhG) from *D. desulfuricans* with known sequences from the MopB superfamily (Wells et al., 2023). Subsequently, these sequences were divided into cytoplasmic Fdhs and FdhG groups, and separate phylogenetic trees were inferred for each.

## 3 Results

### 3.1 Presence or absence of carbon fixation pathway genes

The presence or absence of all the 39 genes for enzymes involved in the WL and rGly pathways and subsequent amino acid synthesis was determined in 54 representative species in the phylum TDB, which is classified into clade A to E (Fig. 2, Supplemental Table S4). Literature search revealed at least 15 out of the 54 species can grow chemolithoautotrophically and they were distributed across clades A, B, C and D (Sánchez-Andrea et al., 2020; Pikuta et al., 2003; Krukenberg et al., 2016; Finster et al., 1998; Moussard et al., 2004; Alain et al., 2010; Lai et al., 2016; Kojima et al., 2016; Hamilton-Brehm et al., 2013; Mardanov et al., 2016; Frolova et al., 2018; Slobodkina et al., 2017; Slobodkin et al., 2013; Cravo-Laureau et al., 2004; DeWeerd et al., 1990). In the 54 representative species, 50 and 44 species possessed the homologs of cytoplasmic Fdhs in *M. thermoacetica* and FdhG in *D. desulfuricans*, respectively (Supplemental Table S4-C). As the results, all the analyzed species was found to have at least one FDHα candidate genes with only one exception of an uncultured *Candidatus* Dadabacteria bacterium (Fig.2). Furthermore, 41 species possessed both cytoplasmic Fdhs and FdhG homologs and the 41 species were distributed in all five clades (A-E) of the phylum TDB. This suggests that the common ancestor of the phylum TDB already possessed genes for both cytoplasmic Fdhs and FdhG.

**Fig. 2.**
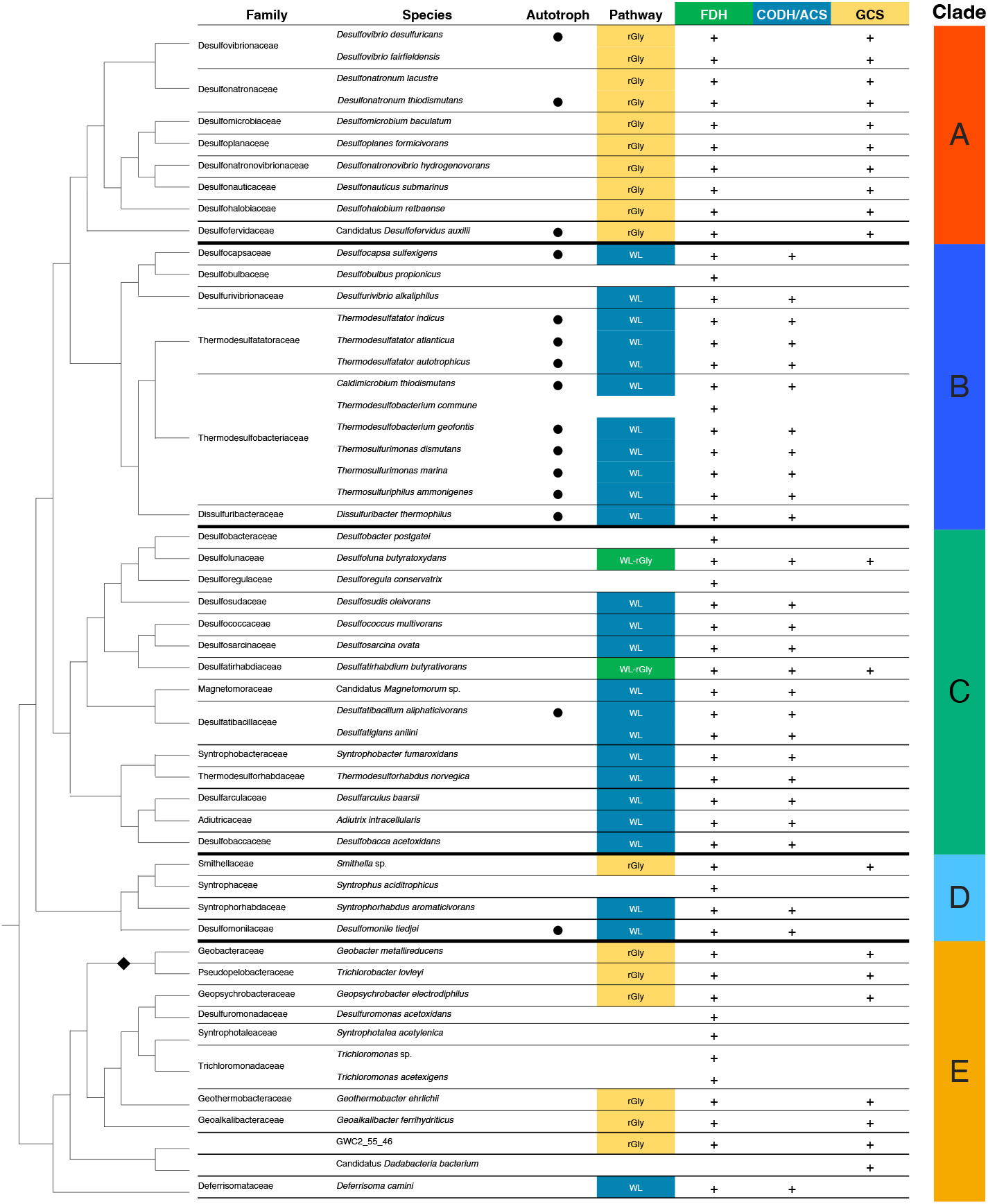
Presence or absence of enzymes for the WL and rGly pathways in Thermodesulfobacteriota species. The tree on the left shows a phylogenetic tree of Thermodesulfobacteriota species, modified from Waite et al.. The order Geobacterales in clade E is indicated by a black diamond (◆). Autotrophic species are indicated by black circles (●). The presence or absence of each enzyme is based on predicted proteomes. FDH is considered to be present (+) when a species encodes either the homolog of cytoplasmic Fdh or FdhG. CODH/ACS is considered to be present if all five subunits of the CODH/ACS complex are encoded. GCS is considered present when all five proteins of the glycine cleavage system are encoded. Pathway: When CODH/ACS and GCS are present together with FDH, the species is estimated to have the WL and rGly pathways, respectively. Clade: the phylum Thermodesulfobacteriota were divided into five monophyletic clades in this study.

The complete gene set for the CODH/ACS complex was identified in 27 species belonging to clades B, C, D, and E, including *T. indicus*, which actually operates the WL pathway for CO_2_ fixation and belongs to clade B (Fig. 2). The complete gene set for the GCS complex was identified in 21 species in clades A, C, D, and E including *D. desulfuricans*, which possesses the rGly pathway and belongs to clade A (Fig. 2). Except for the uncultured *Candidatus* Dadabacteria bacterium, the organisms possessing the gene set for either the CODH/ACS or GCS complex also had FDH genes, suggesting that they may possess the WL or rGly pathway. In particular, species reported to grow autotrophically have high possibility of fixing CO_2_ via either the WL or rGly pathway. All the ten species analyzed in clade A had the gene set for the rGly pathway, whereas none of them had the WL pathway gene set. Conversely, in clade B, 11 out of 13 species conserved the WL pathway gene set, and no species were found to have the rGly pathway gene set (Fig. 2). In contrast to mutually exclusive distribution of the rGly and WL pathways in clade A and B, respectively, both the pathways were estimated to exist in more deeply branching clades C, D, and E compared to clades A and B. Notably, *Desulfoluna butyratoxydans* and *Desulfatirhabdium butyrativorans* in clade C possessed both the gene sets for the WL and rGly pathways. These interspecies distribution of gene sets for the WL and rGly pathways suggested that both the gene sets may have been present in the common ancestor of the phylum TDB.

### 3.2 Phylogenetic analyses of the enzymes for the WL and rGly pathway

If the common ancestor of the phylum TDB possessed the gene sets for both the WL and rGly pathways and these genes were vertically inherited, the phylogeny of these genes would be consistent with the phylogeny of species and exhibited the monophyly in the phylum TDB. Therefore, we conducted phylogenetic analyses focusing on the enzyme genes constituting these pathways to determine whether each enzyme gene was vertically inherited within this phylum or secondarily acquired through horizontal gene transfer (HGT).

A total of 28 phylogenetic trees of the enzymes related to the WL pathway, rGly pathway and subsequent amino acid synthesis were inferred (Figures 3-5 and Supplementary Figures S3-S9). The following section focuses on FDH, CODH/ACS complex, and GCS complex and provides detailed discussions. These three components play particularly important roles: FDH catalyzes the initial step of both the WL and rGly pathways and CODH/ACS and GCS complexes are directly involved in CO_2_ fixation in each pathway.

**Fig. 3.**
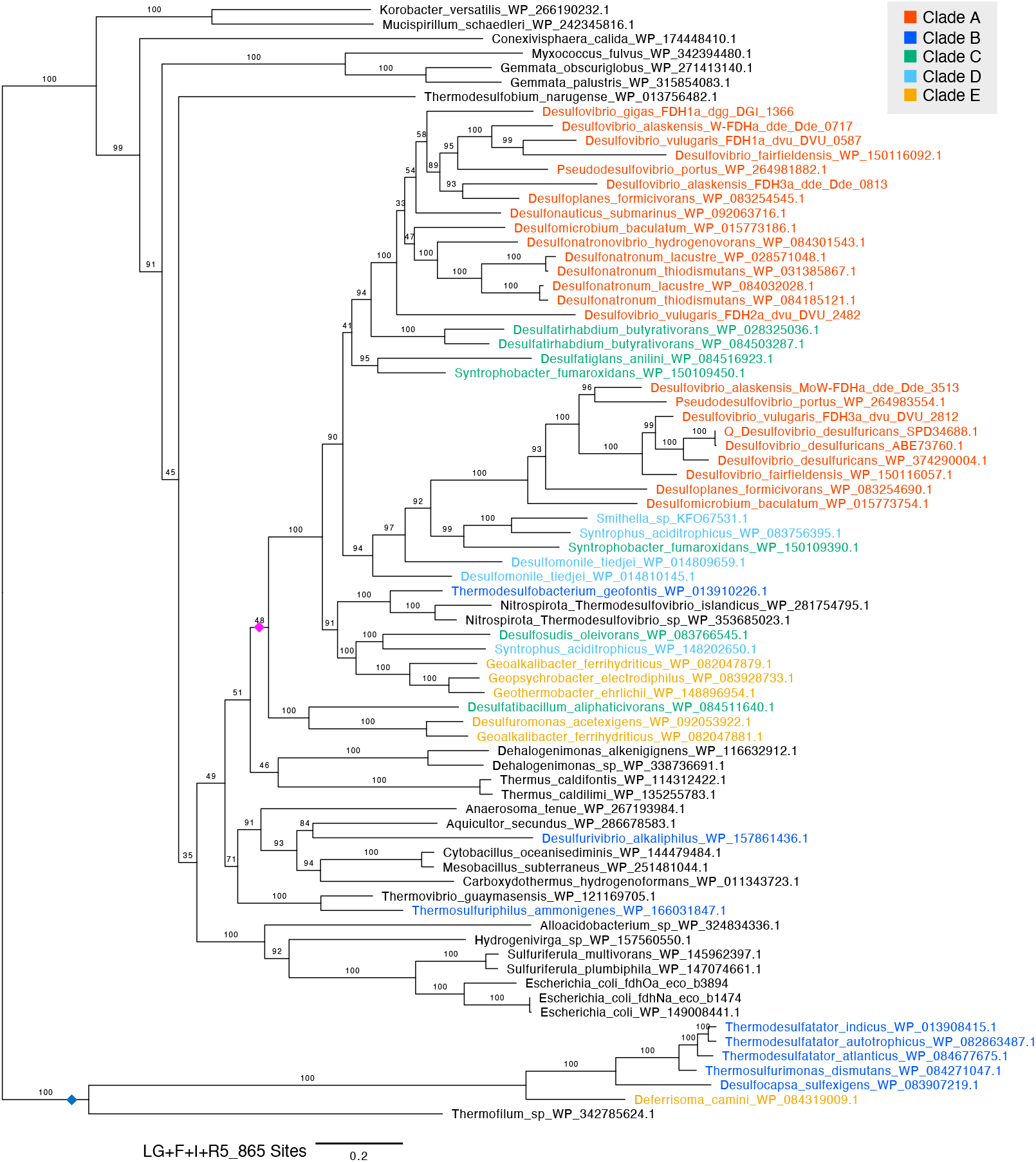
Maximum likelihood phylogenetic tree of FdhG. Bootstrap values were calculated using the ultrafast bootstrap method. The evolutionary model, scale bar and number of sites are shown in the figure. Sequence in the phylum TDB are highlighted with colors. A pink diamond indicates a clade composed of sequences from TDB species. A blue diamond indicates clade B and E sequences that were positioned outside the TDB largest clade.

### 3.3 Phylogenetic analysis of FDHα

In the phylogenetic tree of cytoplasmic Fdh, the sequences from the phylum TDB did not form a monophyletic group (Fig. S3A). This result did not support the vertical transmission of cytoplasmic Fdh gene from the common ancestor of the phylum TDB. In contrast, FdhG from the phylum TDB formed a monophyletic group (Fig. 3, pink diamond). The branching pattern in this monophyletic group was not fully consistent with the phylogenetic relationships of the host species. For instance, sequences from clade A split into two clades. This phylogenetic incongruence can be explained by gene duplications and subsequent gene losses within the phylum TDB. The presence of sequences derives from the genus *Thermodesulfovibrio*, which belongs to the phylum Nitrospirota, is most probably due to the horizontal gene transfer (HGT) from the phylum TDB to the genus *Thermodesulfovibrio*. Therefore, the phylogenetic tree of FdhG supported the presence of FdhG gene in the common ancestor of the phylum TDB.

Within the monophyletic group of FdhG in TDB shown in pink diamond in Fig 3, only one sequence was included from each clade B species, suggesting that FdhG has been lost or undergone substantial genetic divergence in clade B. Two FdhG candidates in clade B organisms (*Desulfurivibrio alkaliphilus* and *Thermosulfuriphilus ammonigenes*) were sporadically distributed outside this monophyletic clade, suggesting the secondarily acquisition through HGT. Furthermore, several FdhG sequences from the clade B together with one from the clade E formed a long-branching clade (Fig. 3, blue diamond) whose evolutionary origin remains unclear.

### 3.4 Phylogenetic analysis of the CODH/ACS complex

In the phylogenetic tree of AcsA, which encodes CODH, the sequences in TDB clustered into two distinct clades supported by bootstrap values (BPs) over 95 each (Fig. 4, highlighted in gray boxes). These two clades were provisionally designated as group 1 and group 2. Group 1 contained AcsA in TDB clades B, C, D, and E, including the one in *T. indicus* possessing the functional WL pathway. In contrast, group 2 contained sequences from clades A, B, C, and E. Eight TDB species encoded two or more AcsA genes within their genomes and each of them were categorized into group 1 and group 2.

**Fig. 4.**
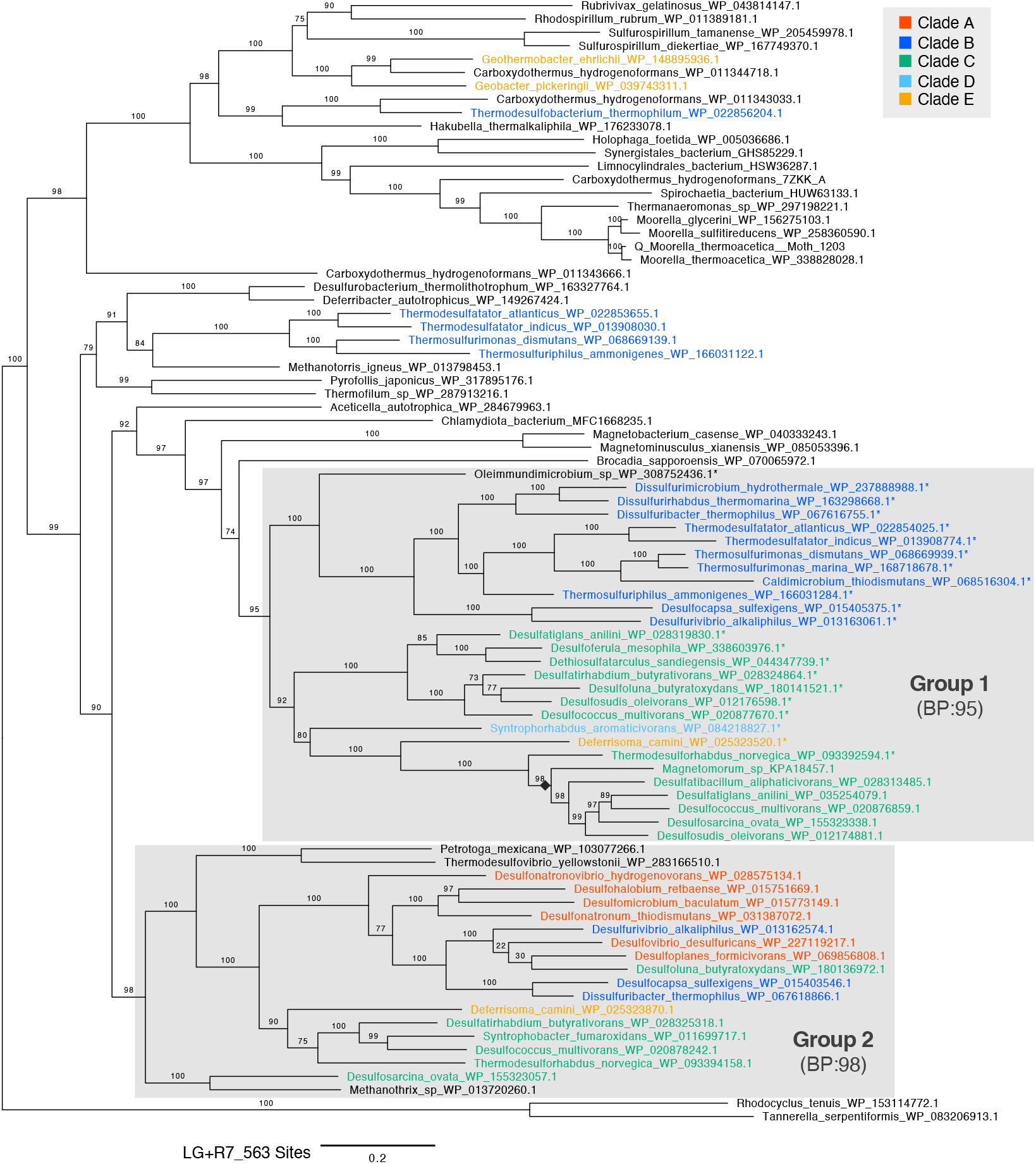
Maximum likelihood phylogenetic tree of AcsA (CODH). Bootstrap values were calculated using the ultrafast bootstrap method. The evolutionary model, scale bar and number of sites are shown in the figure. Sequence in the phylum TDB are highlighted with colors. Asterisks (*) indicate sequences encoded in a cluster with other CODH/ACS complex genes. A black diamond (◆) indicates a clade within group 1 composed of sequences that do not form a gene cluster.

The genomic context analysis of group 1 AcsA revealed that 19 out of 25 TDB-derived AcsA including the one in *T. indicus* formed a cluster with other CODH/ACS complex subunit genes (Fig. 4, highlighted with^*^), supporting its function as a CODH/ACS complex. The other six AcsA without forming the cluster were grouped into one clade (Fig. 4, black diamond). The species with these non-clustered AcsA also retained the previously mentioned AcsA gene that forms the CODH/ACS cluster. Therefore, TDB species with group 1 AcsA may have functional CODH which works in the WL pathway. In contrast, all the 16 AcsA sequences in group 2 derived from the phylum TDB did not form the cluster, suggesting that group 2 AcsA is not involved in acetyl-CoA synthesis (Techtmann et al., 2012). This assumption is supported by group 2 AcsA in *D. desulfuricans*, which has only one AcsA and operates the rGly pathway rather than the WL pathway for CO_2_ fixation.

AcsB subunit encoding ACS which catalyzes acetyl-CoA synthesis with CO, a methyl group of CH_3_- THF, and CoA. All the analyzed AcsB in the phylum TDB, which distributed in all the clades A to E, was clustered into a single clade with a BP support value of 73 (Fig. S4A). Although the phylogeny of AcsB in the phylum TDB was not fully consistent with the phylogeny of the host species, sequences in the clades B and C species formed distinct monophyletic groups supported by a BP of 100. Sequences from Chloroflexota (*Dehalogenimonas formicexedens*) and Planctomycetota (*Kuenenia* sp. and *Scalindua japonica*) were most probably the results of HGT from the phylum TDB to these organisms outside the phylum TDB. Therefore, the phylogenetic analysis supported the hypothesis that the common ancestor of the phylum TDB already possessed the AcsB gene. Phylogenetic analyses also indicate that the AcsC and AcsD genes, which encode subunits involved in methyltransferase activity, were likely present in the common ancestor of the phylum TDB. This is because AcsC and AcsD in the phylum TDB formed monophyletic clades with BPs over 90, respectively (Figs. S4C, D). Similar to AcsB, the AcsC and AcsD clades also contained sequences from the Chloroflexota (*Dehalococcoides* sp.) suggesting that gene cluster-level HGT of the CODH/ACS complex may have occurred from the phylum TDB to the Chloroflexota.

AcsE in the phylum TDB were dispersed across the phylogenetic tree and did not form a monophyletic group (Fig. S4D). Therefore, we could not estimate the origin of AcsE in TDB, which encodes a bacterial-specific subunit involved in THF-corrinoid methyltransferase activity, from the phylogenetic analysis alone.

### 3.5 Phylogenetic analysis of GCS complex

GCSPα in the phylum TDB were divided into two clades provisionally designated as group 1 and 2, supported by a BP of 100 and 94, respectively (Fig. 5A). Group 1 GCSPα was comprised of sequences from clades B, C, D, and E and formed a sister group with GCSPα from the phylum Myxococcota, the closest relative to the phylum TDB. In contrast, group 2 GCSPα included sequences from the species belonging to clade A and the species in the order Geobacterales in clade E. This group 2 also contained sequences from outside the phylum TDB, specifically from Epsilonproteobacteria and the bacterial “PVC group” (Coleman et al., 2021)(Fig. S2). No species possessed both group 1 and group 2 GCSPα genes.

**Fig. 5.**
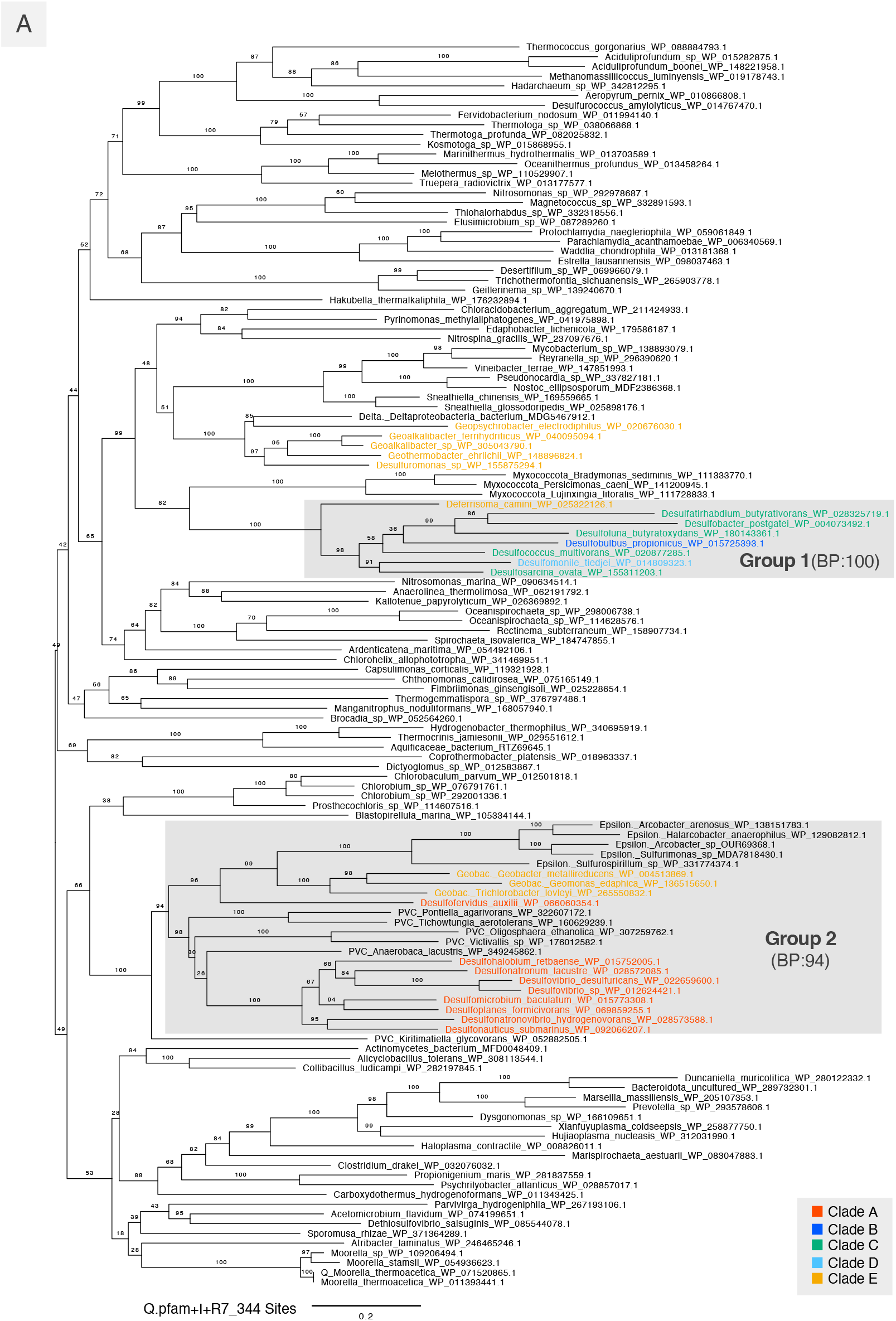
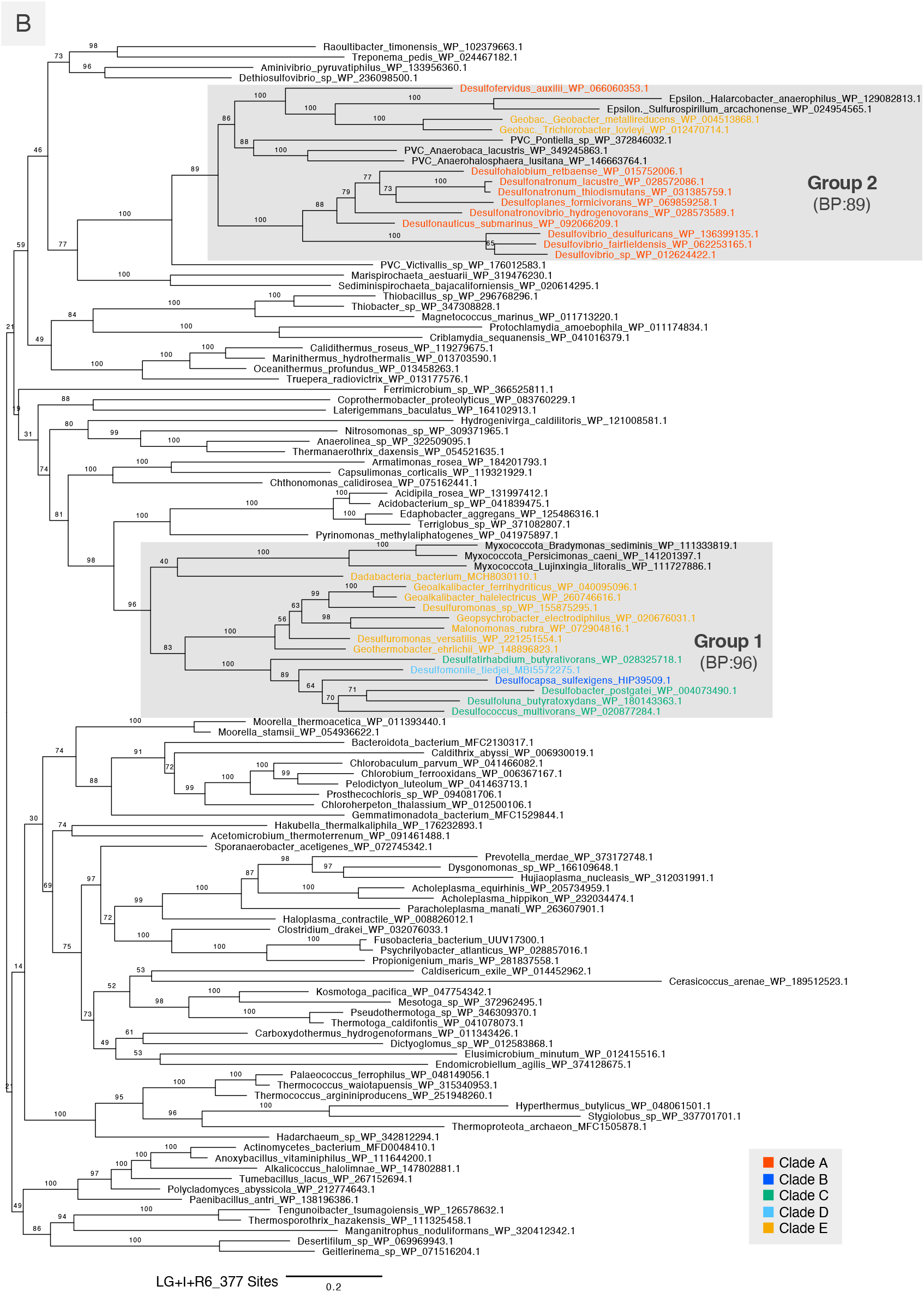
Maximum likelihood phylogenetic tree of GCSP α-subunit (A) and β-subunit (B). Bootstrap values were calculated using the ultrafast bootstrap method. The evolutionary model, scale bar and number of sites are shown in the figure. Sequence in the phylum TDB are highlighted with colors.

Similar to GCSPα, the phylogenetic analyses of GCSPβ and GCST clearly divided the TDB sequences into two groups. Specifically, group 1 consisted of sequences from the TDB clades B, C, D, and E excluding the order Geobacterales (Fig. 5B, Fig. S8C), while group 2 consisted of sequences from clade A and the order Geobacterales in clade E, along with sequences from Epsilonproteobacteria and the PVC group.

GCSPα, GCSPβ and GCST categorized into group 1 based on the phylogenetic trees suggested to be vertically inherited from the common ancestor of the phylum TDB because the sequences from clades B to E formed a monophyletic group although the sequences in the close relatives in the phylum Myxococcota. In contrast, group 2 GCSPα, GCSPβ, and GCST were nested within the PVC group lineage, suggesting that the genes encoding these three subunits were most likely horizontally transferred from the PVC group to clade A and the order Geobacterales.

In contrast to the case of GCSPα, GCSPβ, and GCST, GCSL and GCSH in the phylum TDB were widely scattered across the phylogenetic trees and did not form monophyletic groups. Therefore, we could not estimate the origin of these proteins.

## 4 Discussion

In this study, we investigated the origin and evolutionary history of the WL and rGly pathways in the phylum TDB by analyzing gene presence/absence and phylogeny of the corresponding enzymes. As a result, we succeeded in inferring the gene sets of the common ancestor of the TDB phylum, as well as the subsequent processes of gene gain and loss as described below.

The common ancestor of the phylum TDB is suggested to have possessed the genes for CODH/ACS complex. Four out of five CODH/ACS complex subunits, AcsA, B, C, and D, are inferred to have existed in the common ancestor of the TDB phylum based on gene presence/absence and phylogenetic analyses. Although our phylogenetic analysis did not reveal the origin of AcsE, this protein is also expected to have existed in the common ancestor of the phylum TDB. This is because AcsE gene forms a cluster with other CODH/ACS complex genes and is therfore most likely vertically inherited as a cluster.

The common ancestor of the phylum TDB may also have possessed the genes for GCS complex. Both phylogenetic and gene presence/absence analyses supported the vertical inheritance of GCSPα, GCSPβ, and GCST to clade B, C, D and E from the common ancestor of TDB. In contrast, phylogenetic analyses did not clarify the origin of GCSL and GCSH in the phylum TDB although they are widely distributed in all the clades (Supplementary Table S4-D). This could be explained by rapid sequence evolution caused by low functional constraints or replacement by functionally equivalent proteins because GCSH is a non-catalytic carrier protein and GCSL is an FAD dependent oxidoreductase that re-oxidizes the H-protein (Kikuchi et al., 2008; Kikuchi, 1973). Therefore, we estimated that the common ancestor of TDB may also have possessed the genes for GCSL and GCSH.

The common ancestor of the phylum TDB is also expected to have FdhG, a catalytic unit of FDH, indicating that it may have all the key enzyme genes for the WL and rGly pathways. Furthermore, presence/absence analysis also detected all the genes other than those for the key enzymes for the WL and rGly pathways in the phylum TDB (Supplementary Table S4-D). Namely, enzymes involved in the methyl branch which are shared between the WL and rGly pathways and an enzyme catalyzing the subsequent reaction in the WL pathway (Fig. S1) were identified in all the clades. These findings support the possibility that the common ancestor of the phylum TDB possessed the gene set for the WL and rGly pathways.

Based on the phylogeny of enzymes, the following evolutionary scenario is proposed in the phylum TDB (Fig. 6). The gene set for CODH/ACS complex was lost in the common ancestor of clade A and in clade E after the ancestor of *D. camini* was separated from others (Fig. 6, Blue). GCS complex genes were lost in the common ancestor of clades A-B and in the order Geobacterales within clade E (Fig. 6, Yellow). The CODH/ACS complex and/or GCS complex have also been lost in some species in clades B to E multiple times. Moreover, clade A and the order Geobacterales regained group 2 GCSPα, Pβ, and T from PVC group via HGT (Fig. 6, Pink). These findings indicate that individual lineages have formed unique gene repertoires through gene losses and acquisitions via HGT.

**Fig. 6.**
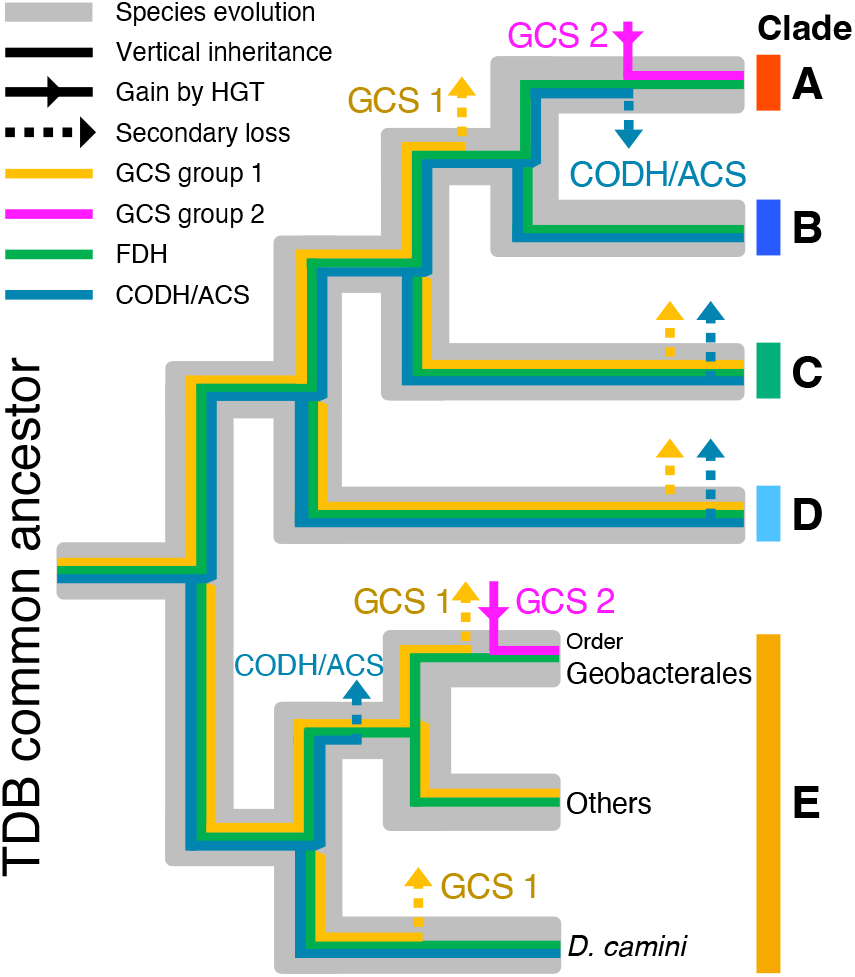
Gain and loss scenarios of gene sets for the WL and rGly pathways in the phylum Thermodesulfobacteriota. The phylogenetic tree of the phylum TDB is shown gray. Presence of absence of FDH, CODH/ACS complex, and GCS complex are shown with color bars explained in the figure.

Although the common ancestor of TDB is expected to have possessed all the enzyme genes for the WL and rGly pathways, it remains unsolved whether these genes have worked for CO_2_ fixation. To estimate their function for autotrophic growth, we analyzed downstream of the carbon fixation pathways. If the common ancestor of the TDB phylum grew autotrophically via the WL pathway, it would have pyruvate ferredoxin oxidoreductase, phosphoserine phosphatase, and serine hydroxymethyltransferase (Fig. 1), because these enzymes are essential to synthesize serine and glycine from acetyl-CoA through pyruvate. As a result, the three enzymatic genes were detected in all clades of the phylum TDB (Table S4-D). Although phylogenetic analyses of these three enzymes did not support their vertical inheritance from the common ancestor of the phylum TDB (Fig. S4-6) plausibly due to frequent HGT and gene duplication events, these were likely present in the common ancestor of TDB. Therefore, the common ancestor of TDB may have been capable of autotrophic growth via the WL pathway.

We also analyzed the five subunits of the glycine reductase complex (Fig. 1), which catalyzes the conversion of glycine to acetyl-CoA via acetyl phosphate and therefore reported to be essential for autotrophic growth via the rGly pathway (Sánchez-Andrea et al., 2020; Song et al., 2020). Neither gene presence/absence nor phylogenetic analysis supported the presence of the glycine reductase complex in the common ancestor. Still, there remains a possibility that glycine produced via the rGly pathway had been converted to serine by SHMT and then to pyruvate. The common ancestor of the phylum TDB might have grown autotrophically by operating both the WL and rGly pathways like *C. drakei* (Song et al., 2020). Therefore TDB might have been capable of autotrophic growth via the rGly pathway.

To date, no species within the phylum TDB has been reported to operate both the WL and rGly pathways. Our presence/absence analysis demonstrates that two species in clade C (*Desulfoluna butyratoxydans* and *Desulfatirhabdium butyrativorans*) possess the key gene sets for both the WL and rGly pathways (Fig. 2), however, both species are known as heterotrophs (Suzuki et al., 2008; Balk et al., 2008). Further studies are needed to address the possibility that they grow autotrophically under unknown conditions and/or these metabolic pathways function as auxiliary pathways for organic compound acquisition or as sink for reducing power (Sánchez-Andrea et al., 2020; Schneeberger, 1999).

LBCA is estimated to have a gene set for the rGly pathway except for FDH (Coleman et al., 2021). However, their study used only cytoplasmic Fdh (K05299, K15022) as the query for FDH based on the information of KEGG pathway map, in which only cytoplasmic Fdh is registered (Aug 2025). Nonetheless, FDH has multiple isomers other than cytoplasmic Fdh, including FdhG that converts CO_2_ to formate in the rGly pathway of *D. desulfuricans* (Wells et al., 2023). Therefore, it remains possible that the LBCA did possess FDH that is not classified as cytoplasmic Fdh, like the case of the common ancestor of the phylum TDB which may have possessed FdhG.

In this study, we focused on a specific phylum and estimated the metabolisms of its last common ancestor. This phylum-specific studies ensures appropriate taxon sampling and enables more accurate inference of ancestral traits than the studies focused on deeper lineage, such as LBCA (Powell and Battistuzzi, 2022; Superson et al., 2019). Accumulation of knowledges based on such clade-specific studies across various phyla will lead to a more precise inference of metabolism in ancestral organisms such as LBCA and LUCA.

## 6 Author Contributions

TW, KK and YC conceived and designed the work and wrote the manuscript. TW performed the presence/absence analyses. TW and KK inferred the phylogenetic trees. TW, KK and YC conducted phylogenetic and evolutionary analyses. All authors reviewed and approved the final version of the manuscript.

## 8 Acknowledgments

We appreciate the fruitful discussions with Dr. Tetsuo Hashimoto regarding the phylogenetic analyses and evolutionary interpretations. We also thank Mr. Kohei Bamba for his assistance with the single-gene phylogenetic analyses. We are grateful to Dr. Ryuhei Nakamura, Dr. Eric Smith and Dr. Kimiho Omae for their helpful discussions.

