## Supplementary_Material for "Molecular evolution of the Wood-Ljungdahl pathway and the reductive glycine pathway in Thermodesulfobacteriota"

### 1 Supplementary Tables

**Supplementary Table 1.** Subunit composition of the CODH/ACS complex.
Presented as a separate MS Excel file.

**Supplementary Table 2.** Queries used for presence/absence analyses.
Presented as a separate MS Excel file.

**Supplementary Table 3.** Queries and summary for phylogenetic analyses.
Presented as a separate MS Excel file.

**Supplementary Table 4.** Raw data for gene presence/absence analyses.
Presented as a separate MS Excel file.

### 2 Supplementary Figures

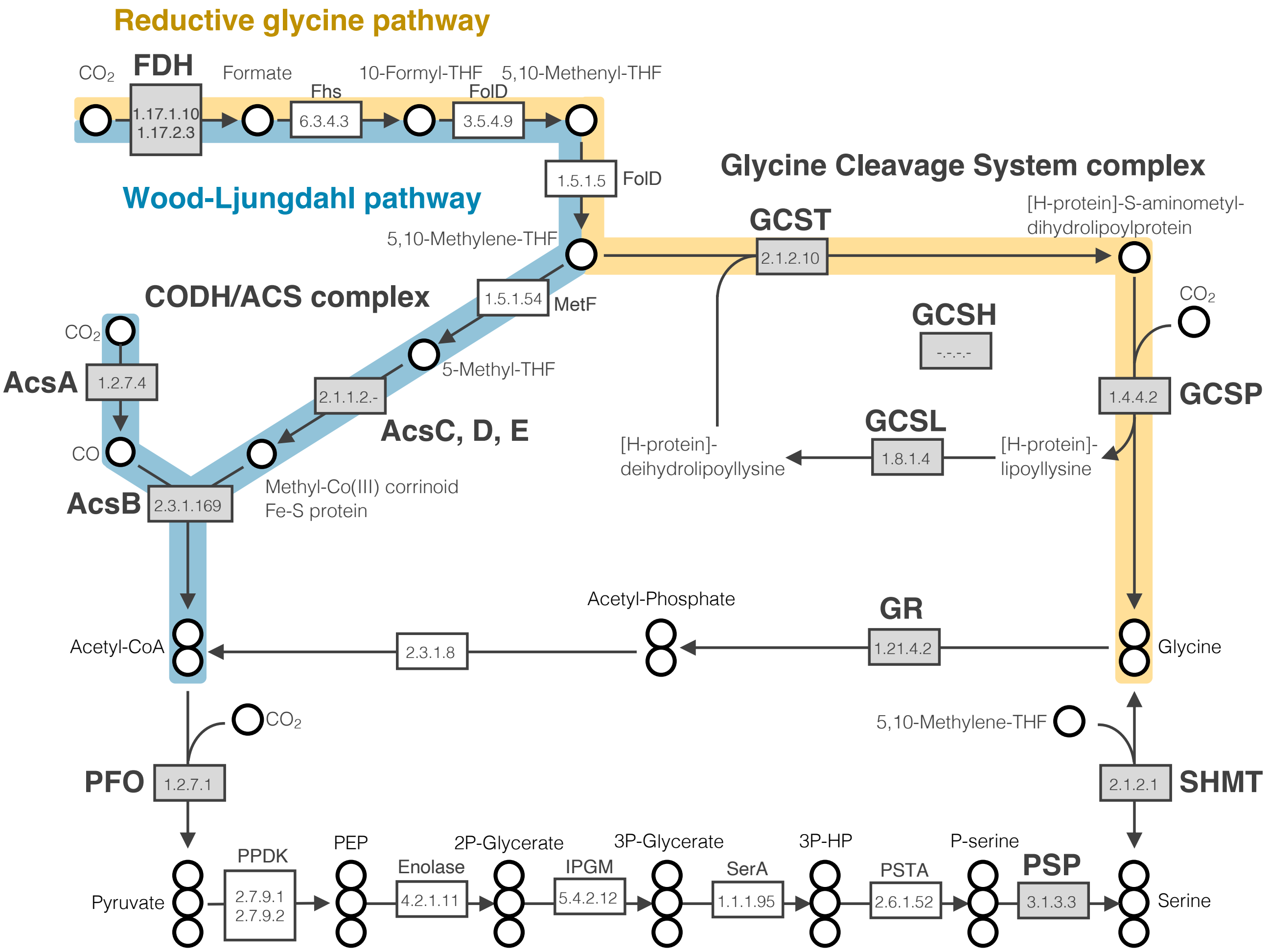


**Supplementary Figure 1. The WL and rGly pathways and related metabolism.** Each arrow represents an enzymatic reaction and box indicates the EC number. Gray boxes highlighted enzymes subjected to phylogenetic analysis. Enzyme names are shown in bold. Circles represent carbon. THF: tetrahydrofolate; FDH: formate dehydrogenase; CODH/ACS: carbon monoxide dehydrogenase/acetyl-CoA synthase; GCS: glycine cleavage system; GCST: glycine cleavage system T-protein; GCSP: glycine cleavage system P-protein; GCSL: glycine cleavage system L-protein; GCSH: glycine cleavage system H-protein; PFO: pyruvate-ferredoxin oxidoreductase; SHMT: serine hydroxymethyltransferase; PSP: phosphoserine phosphatase; GR: glycine reductase; PEP: phosphoenolpyruvate; 2P-Glycerate: 2-phosphoglycerate; 3P-Glycerate: 3-phosphoglycerate; 3P-HP: 3-phosphohydroxypyruvate; P-serine: phosphoserine.

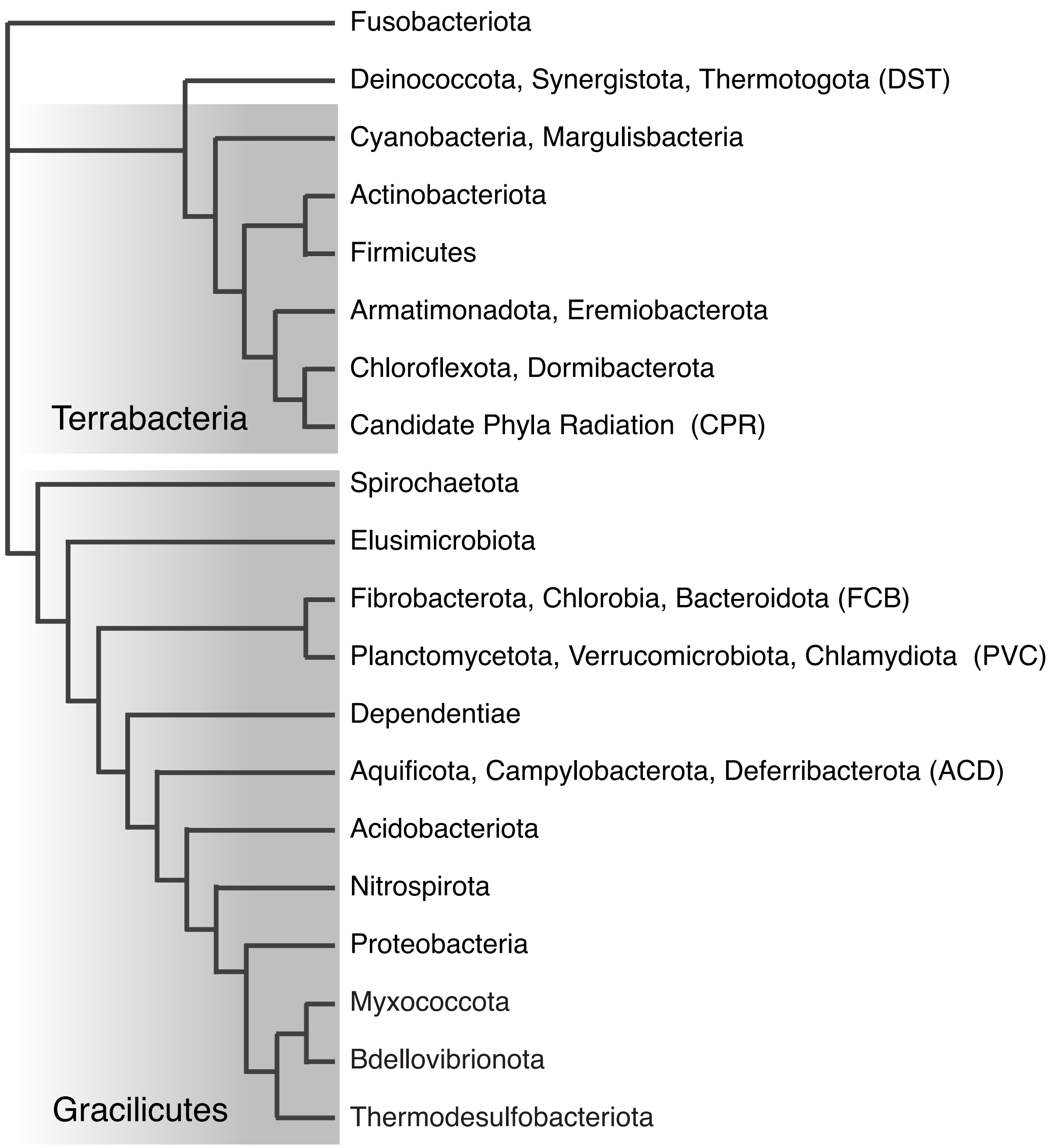


**Supplementary Figure 2. Unrooted phylogenetic tree of the bacterial domain.**
An unrooted phylogenetic tree of the bacterial domain based on Coleman et al. and Waite et al.

【Fig. S3A. Maximum likelihood phylogenetic tree of cytoplasmic Fdh α-subunit.】

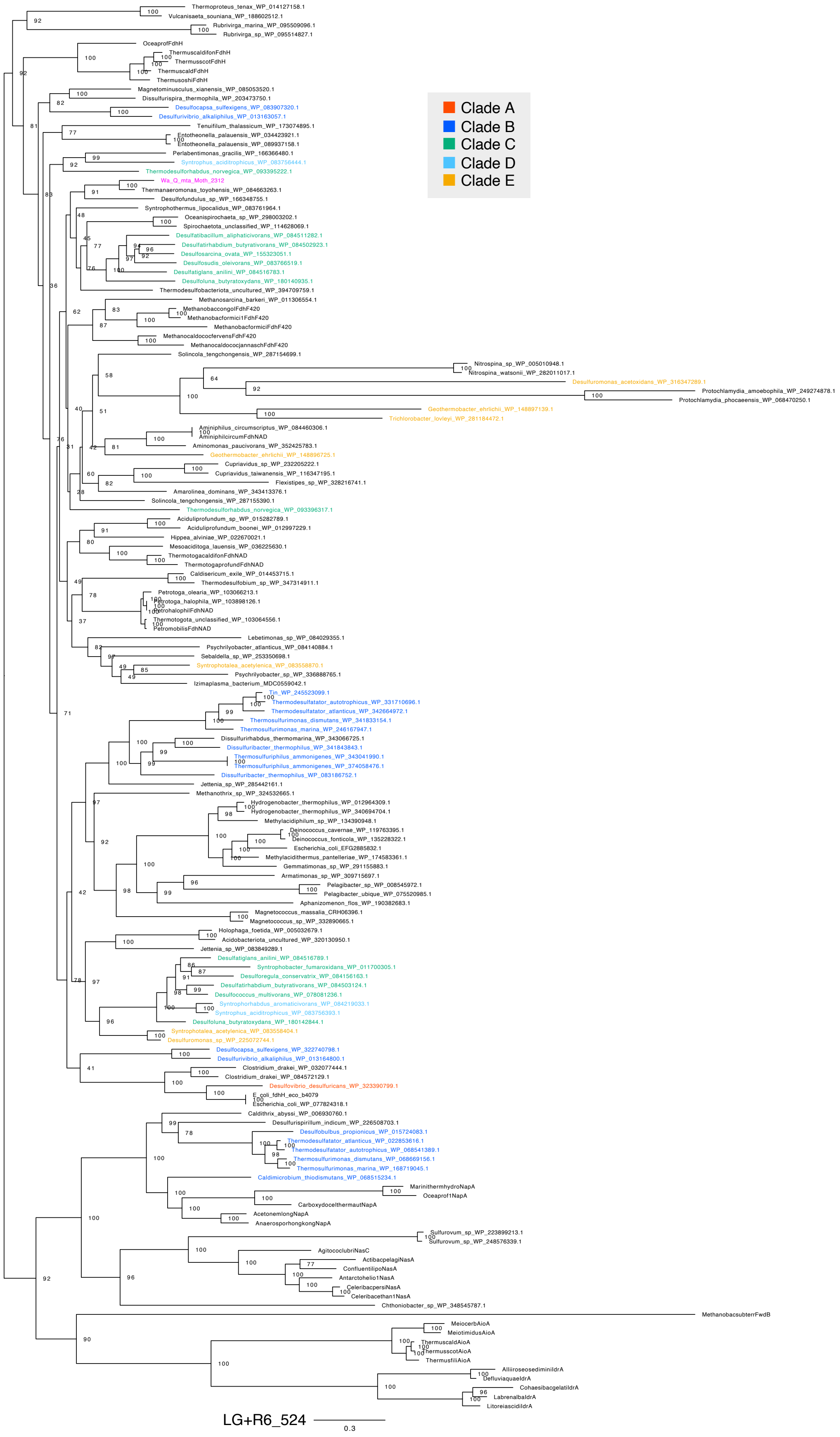

【Fig. S3B. Maximum likelihood phylogenetic tree of cytoplasmic Fdh β-subunit.】

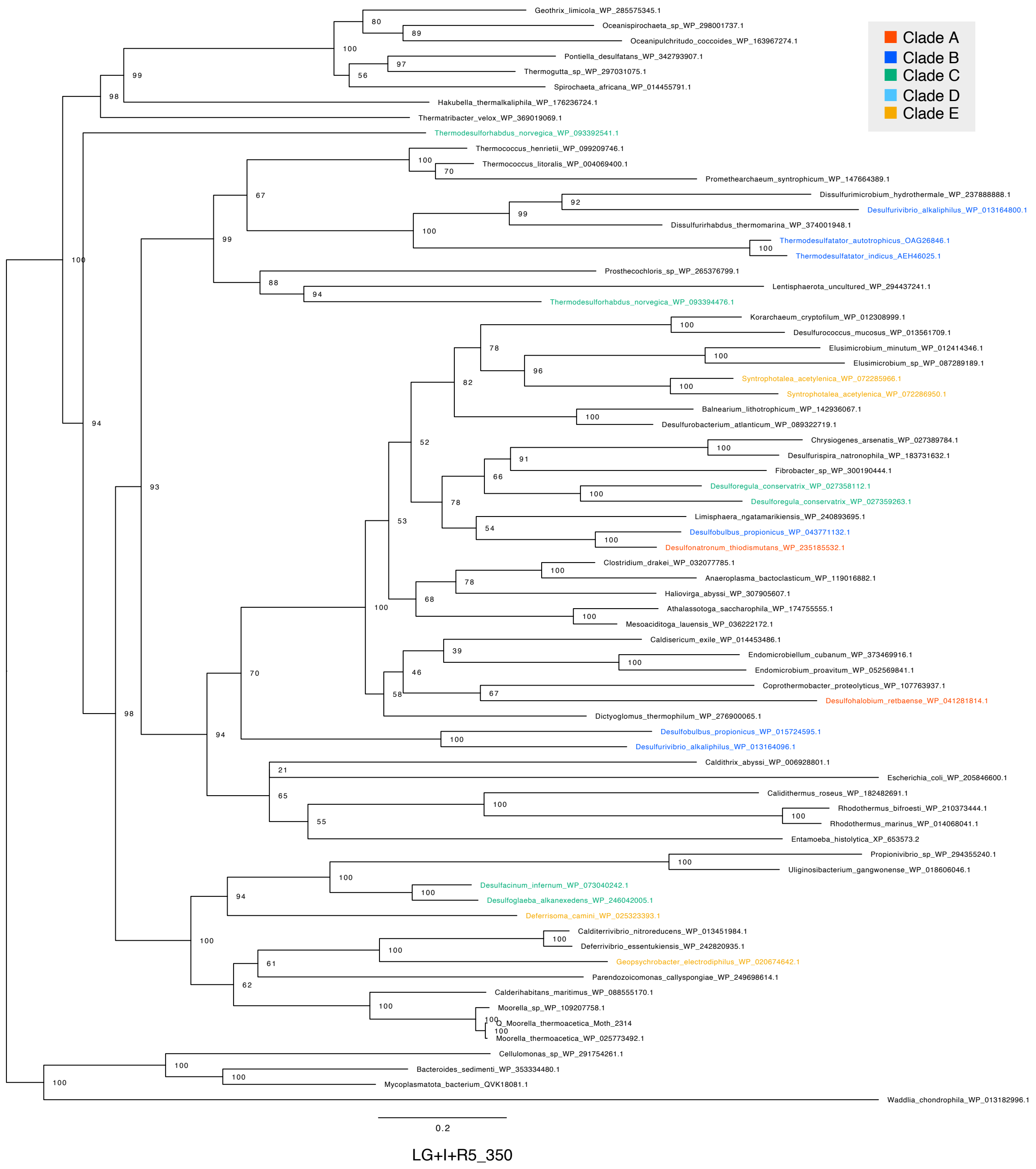

【Fig. S3C. Maximum likelihood phylogenetic tree of FdhG β-subunit.】

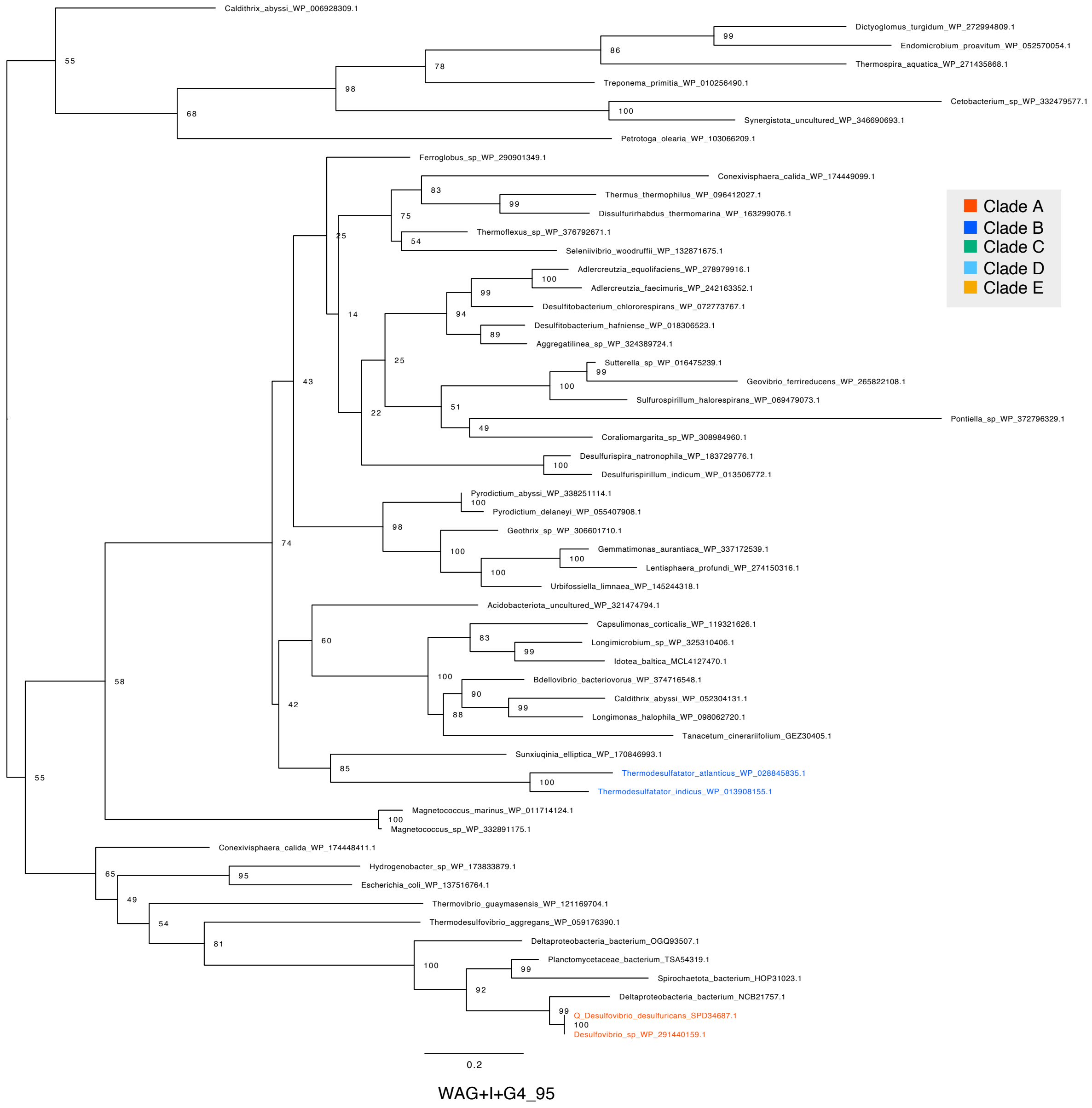

【Fig. S3D. Maximum likelihood phylogenetic tree of FdhG γ-subunit.】

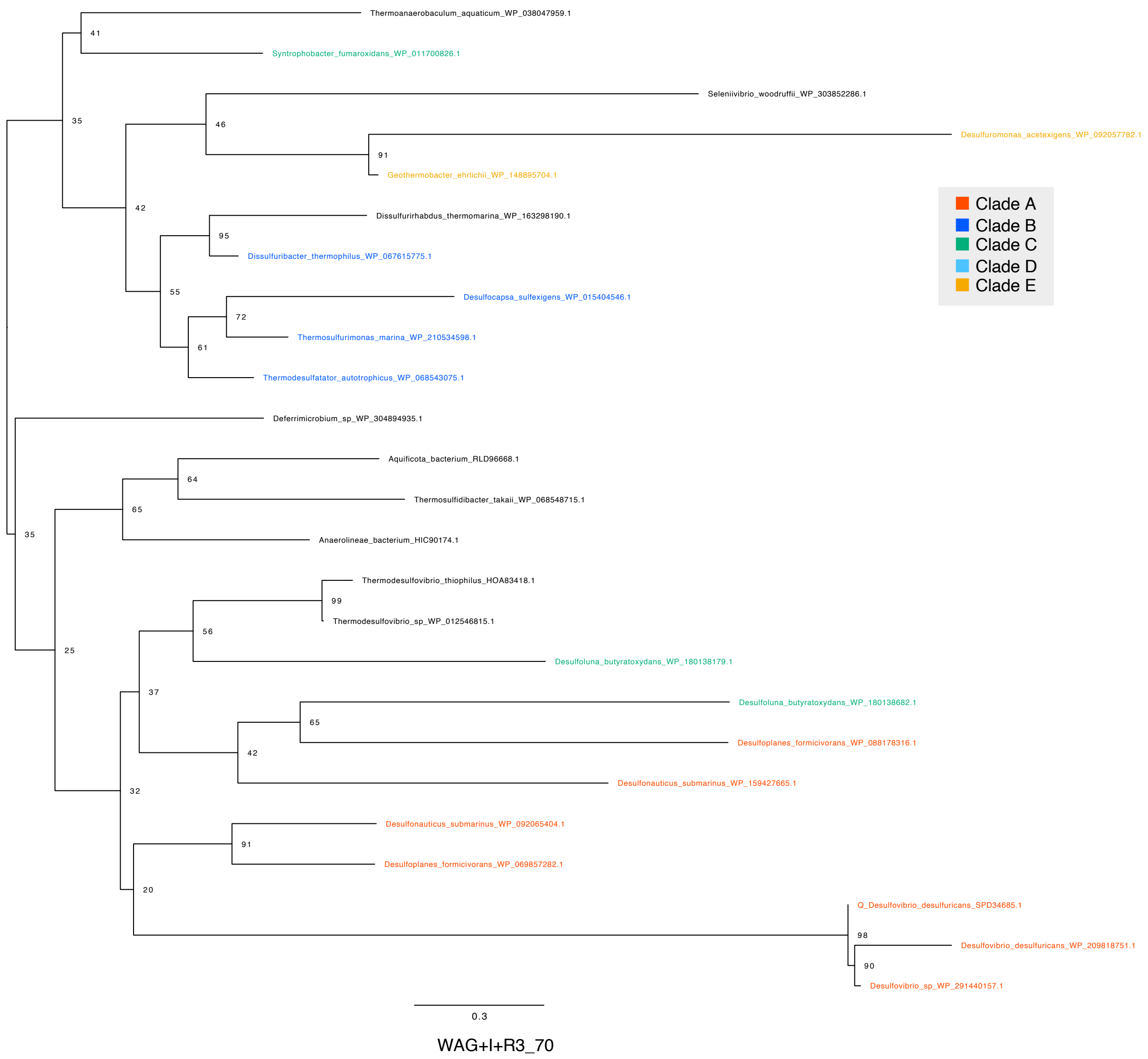


**Supplementary Figure 3. Maximum likelihood phylogenetic tree of FDH.**
(A) Cytoplasmic Fdh α-subunit; (B) Cytoplasmic Fdh β-subunit; (C) FdhG β-subunit; (D) FdhG γ-subunit. Bootstrap values were calculated using the ultrafast bootstrap method. The evolutionary model, scale bar and number of sites are shown in the figure. Sequence in the phylum TDB are highlighted with colors.

【Fig. S4A. Maximum likelihood phylogenetic tree of AcsB.】

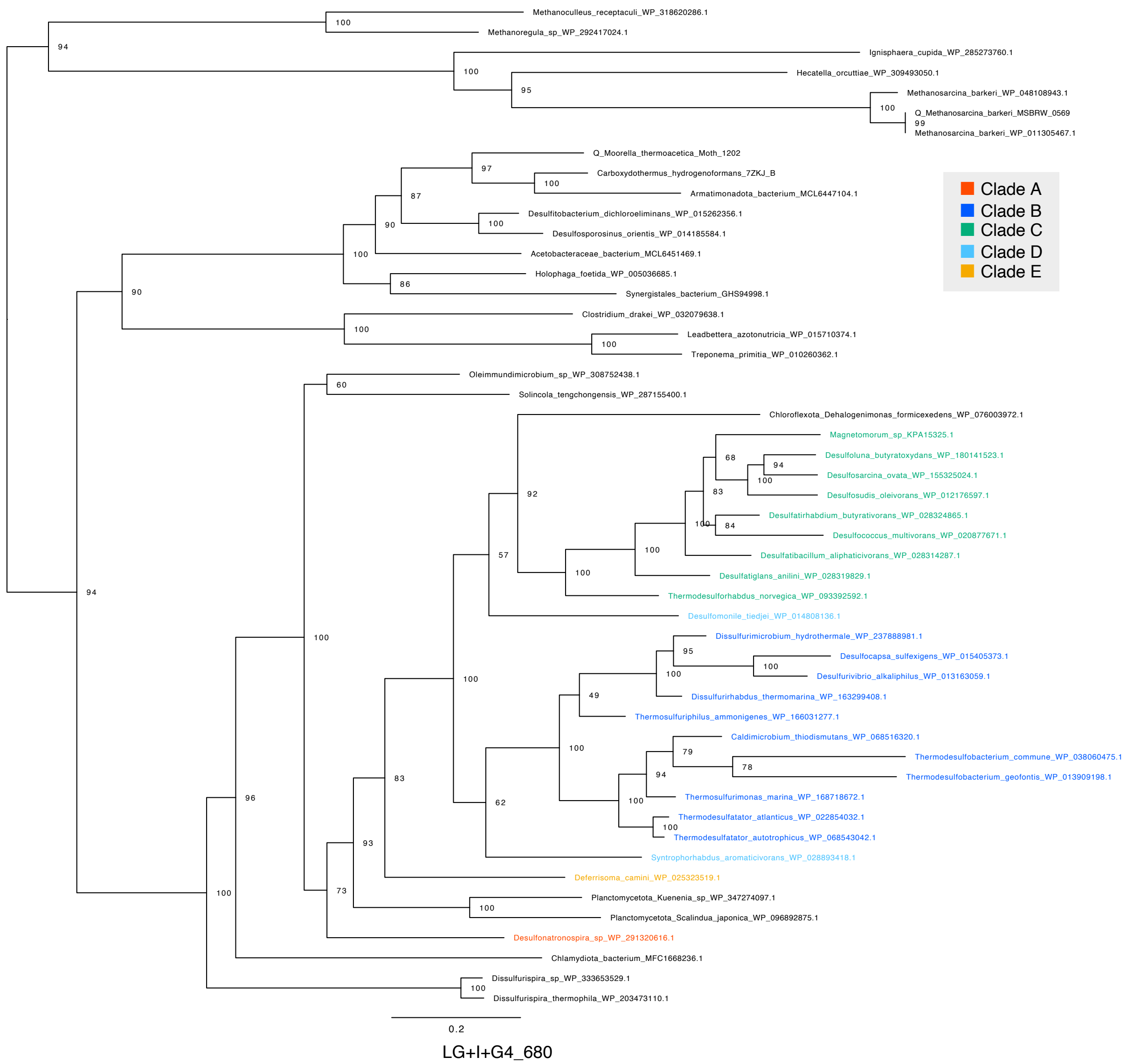

【Fig. S4B. Maximum likelihood phylogenetic tree of AcsC.】

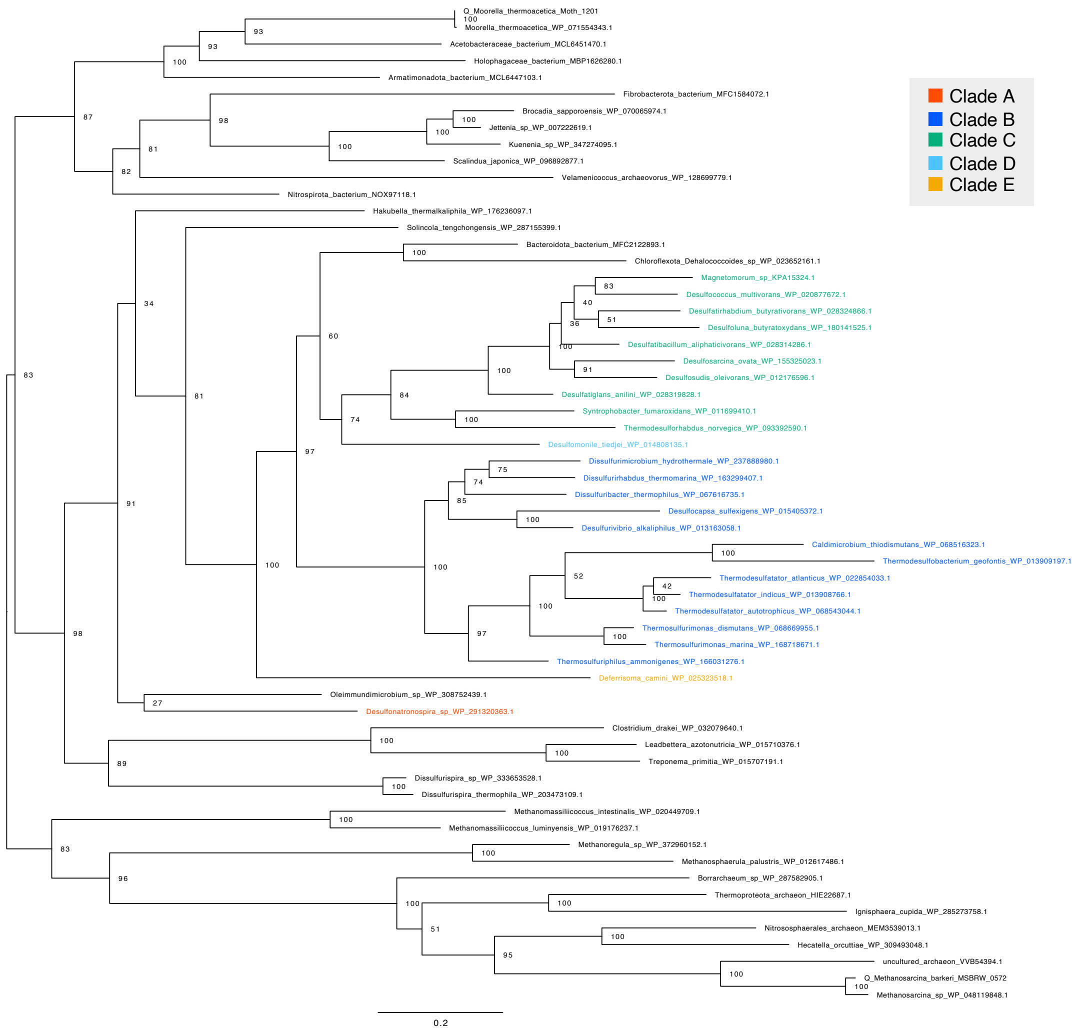


LG+F+I+G4_402

【Fig. S4C. Maximum likelihood phylogenetic tree of AcsD.】

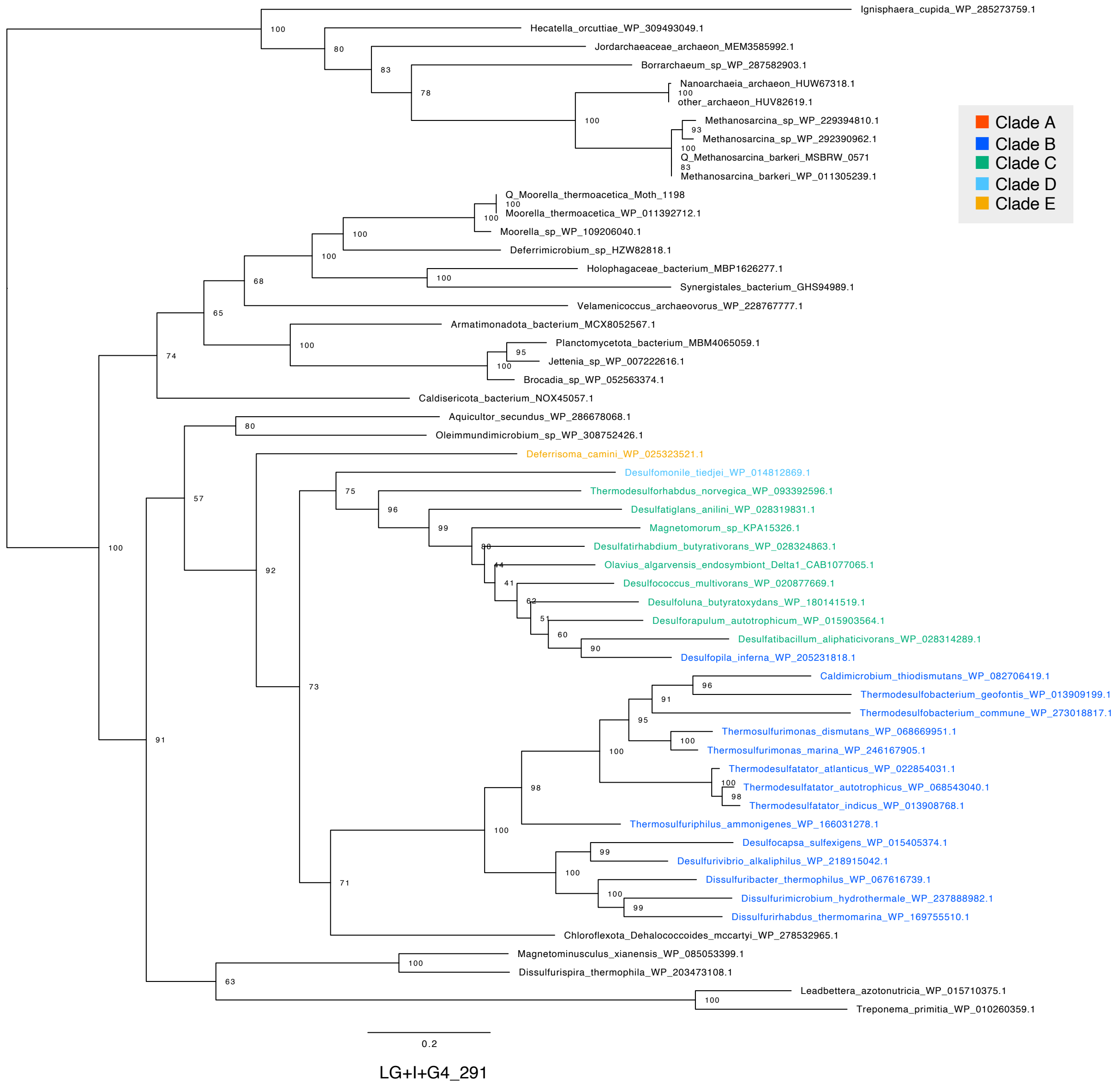

【Fig. S4D. Maximum likelihood phylogenetic tree of AcsE.】

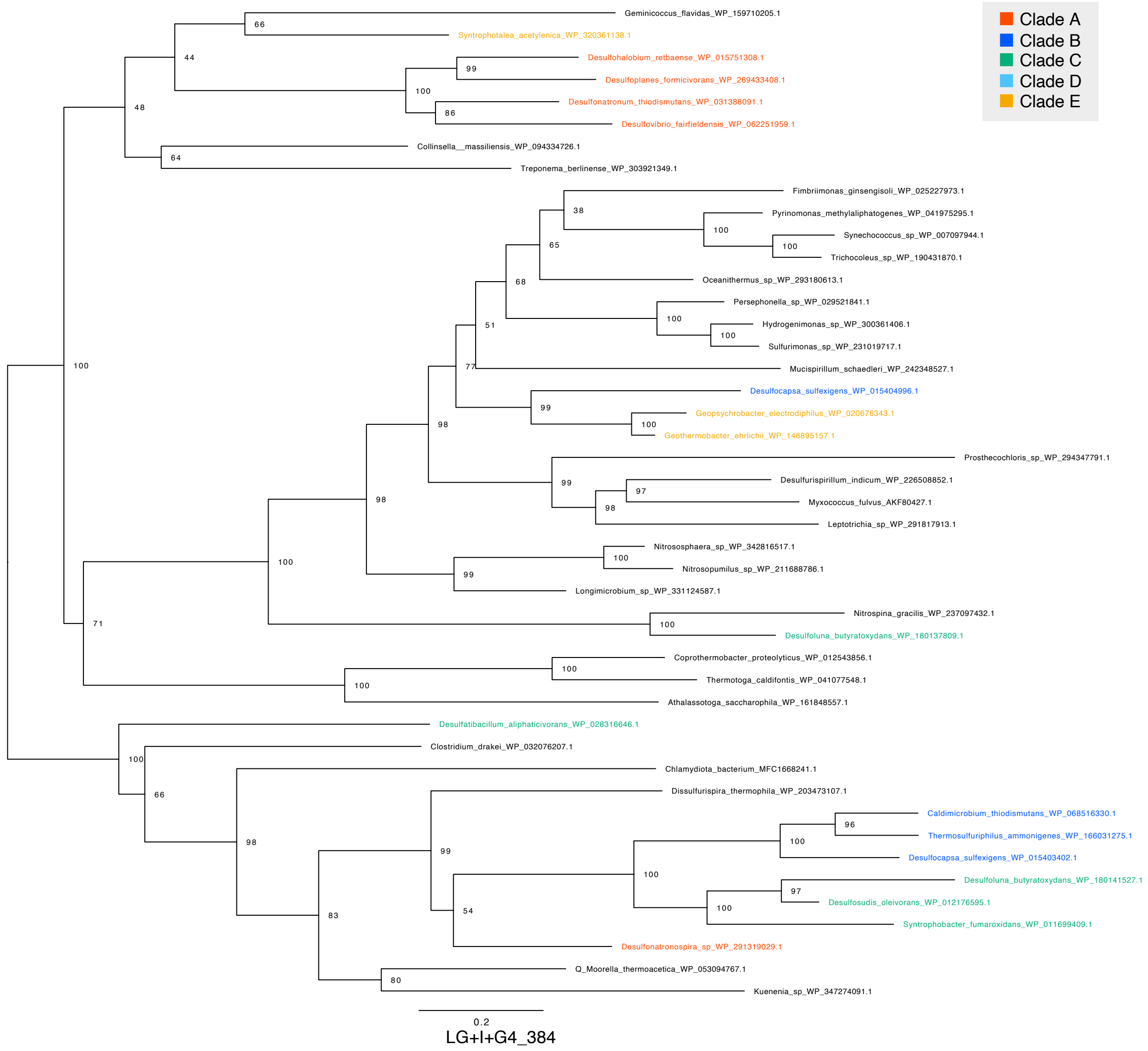


**Supplementary Figure 4. Maximum likelihood phylogenetic tree of CODH/ACS complex.** (A) AcsB; (B) AcsC; (C) AcsD; (D) AcsE. Bootstrap values were calculated using the ultrafast bootstrap method. The evolutionary model, scale bar and number of sites are shown in the figure. Sequence in the phylum TDB are highlighted with colors.

【Fig. S5A. Maximum likelihood phylogenetic tree of PFO (fysed).】

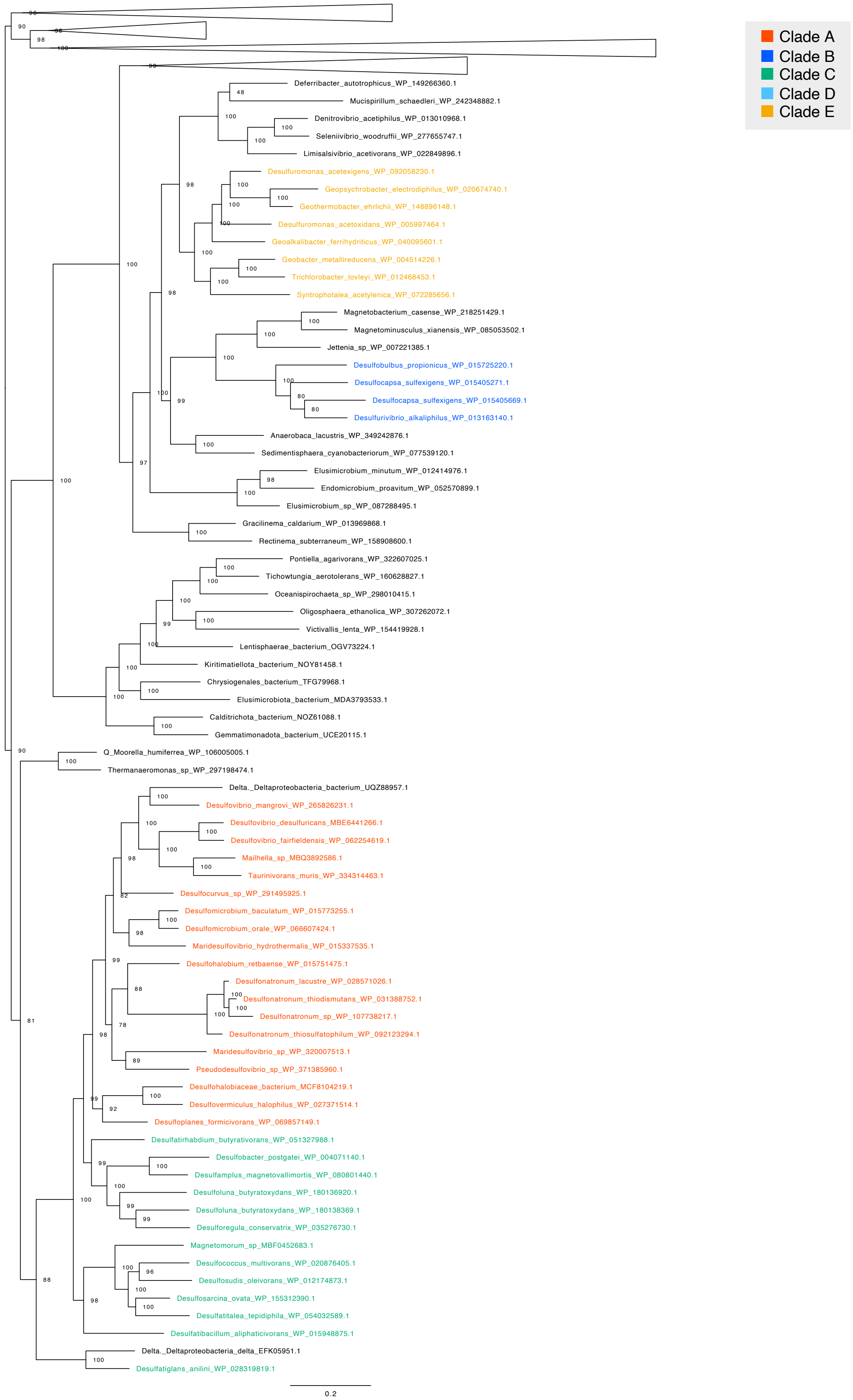


LG+F+I+R10_933

【Fig. S5B. Maximum likelihood phylogenetic tree of PFO α-subunit.】

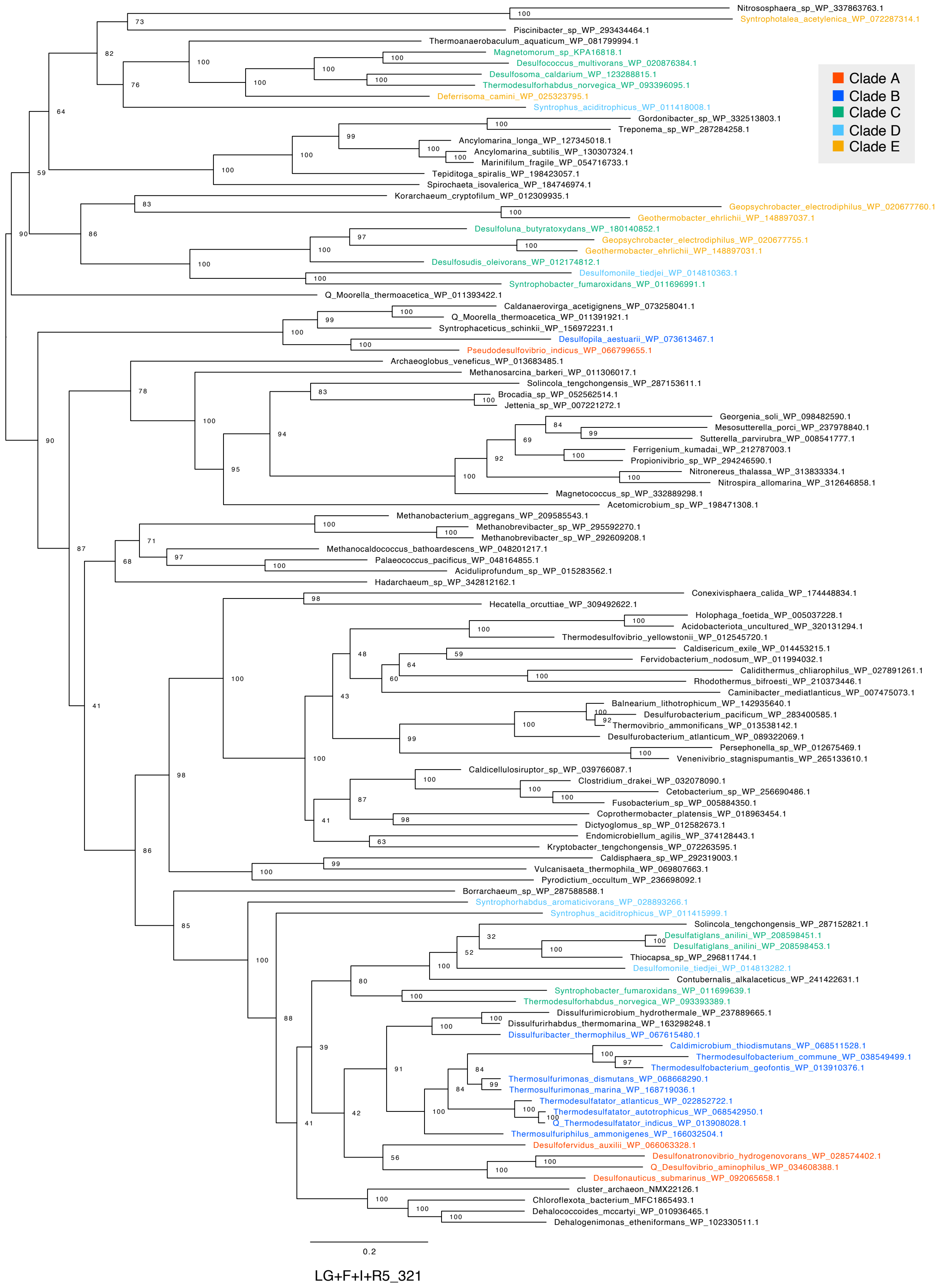

【Fig. S5C. Maximum likelihood phylogenetic tree of PFO β-subunit.】

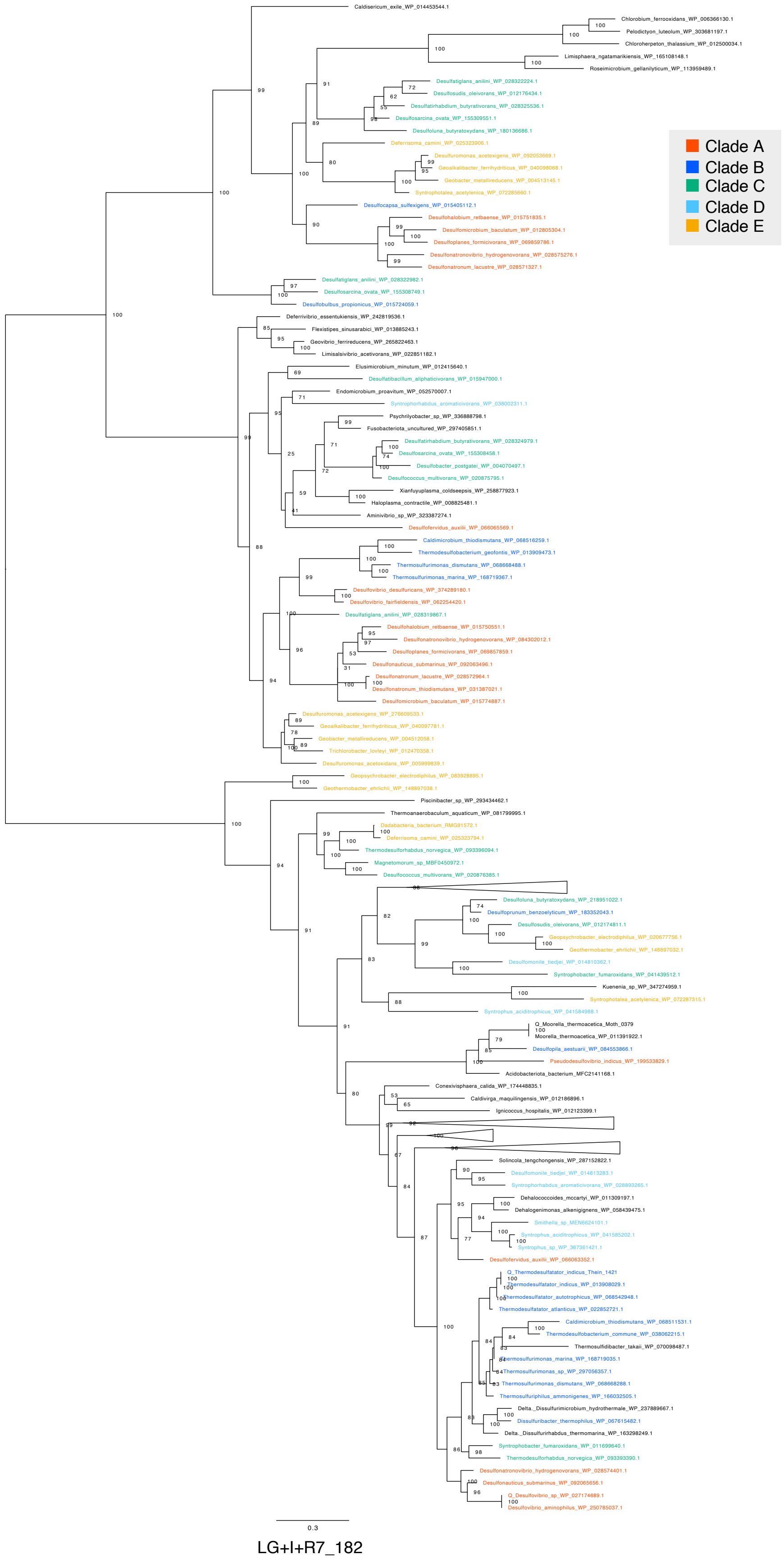

【Fig. S5D. Maximum likelihood phylogenetic tree of PFO γ-subunit.】

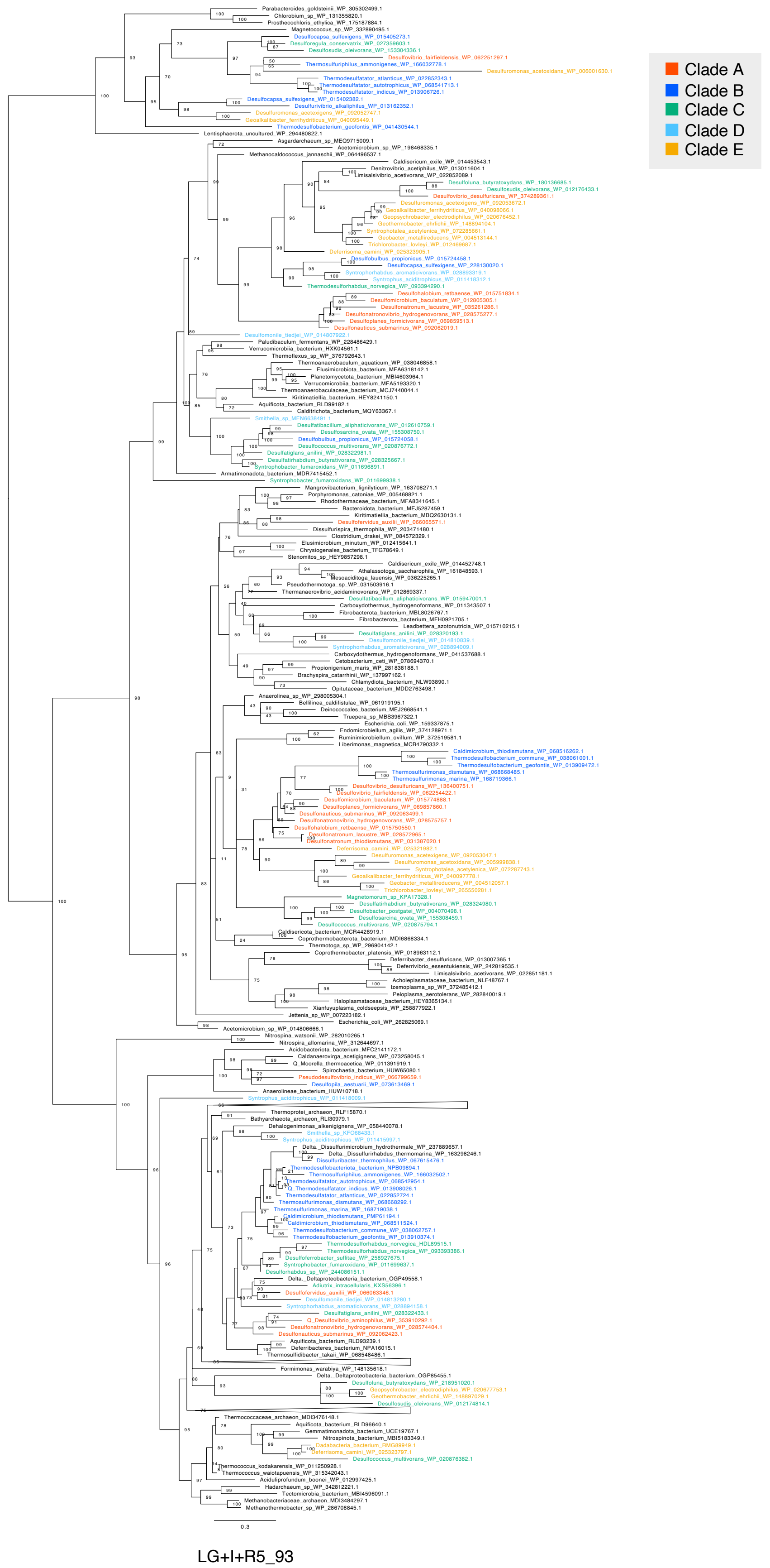

**Supplementary Figure 5. Maximum likelihood phylogenetic tree of PFO.**
(A) PFO (fused); (B) PFO α-subunit; (C) PFO β-subunit; (D) PFO γ-subunit.
Bootstrap values were calculated using the ultrafast bootstrap method. The evolutionary model, scale bar and number of sites are shown in the figure. Sequence in the phylum TDB are highlighted with colors.

【Fig. S6A. Maximum likelihood phylogenetic tree of dPSP1.】

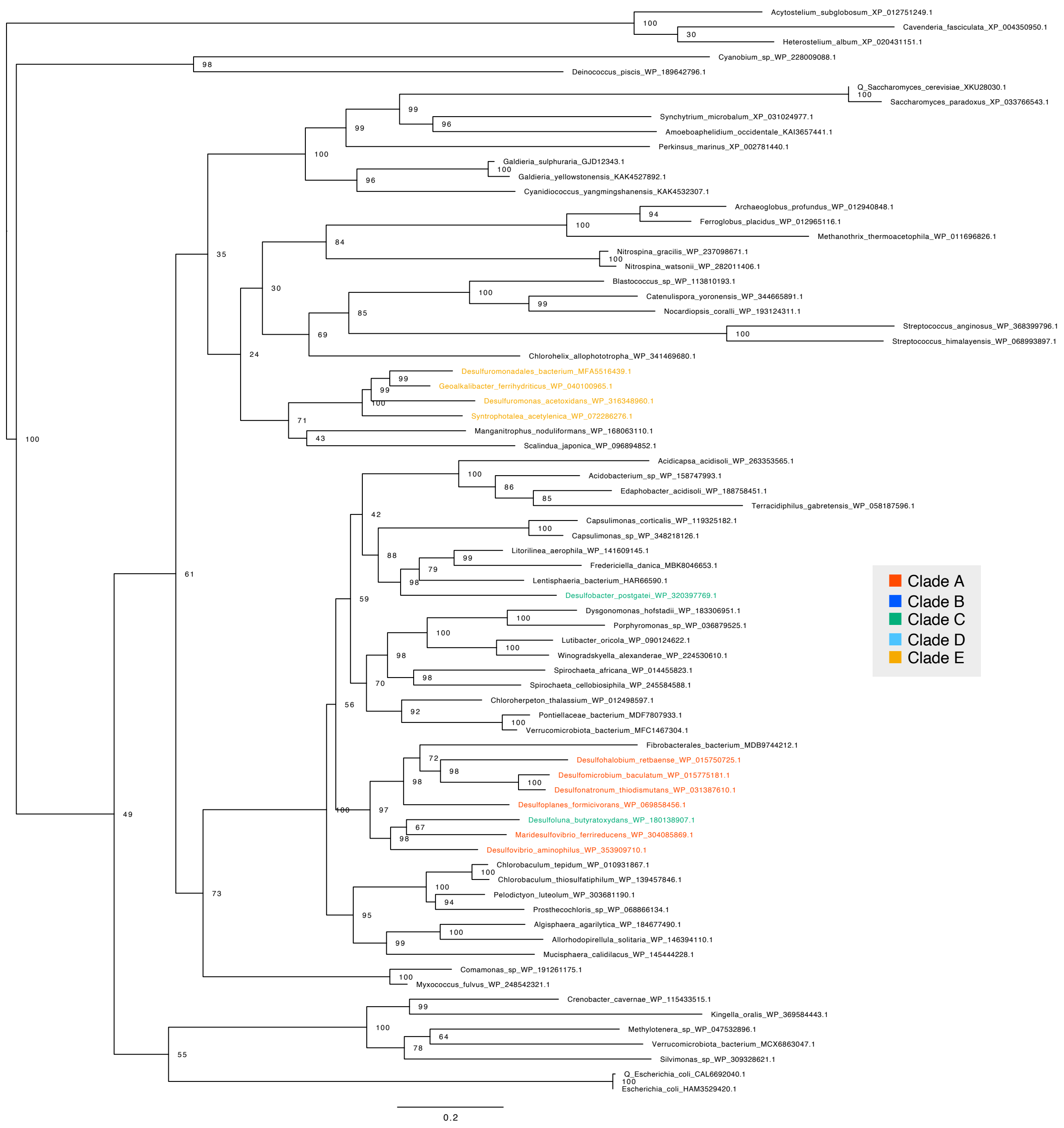


LG+R5_270

【Fig. S6B. Maximum likelihood phylogenetic tree of dPSP2.】

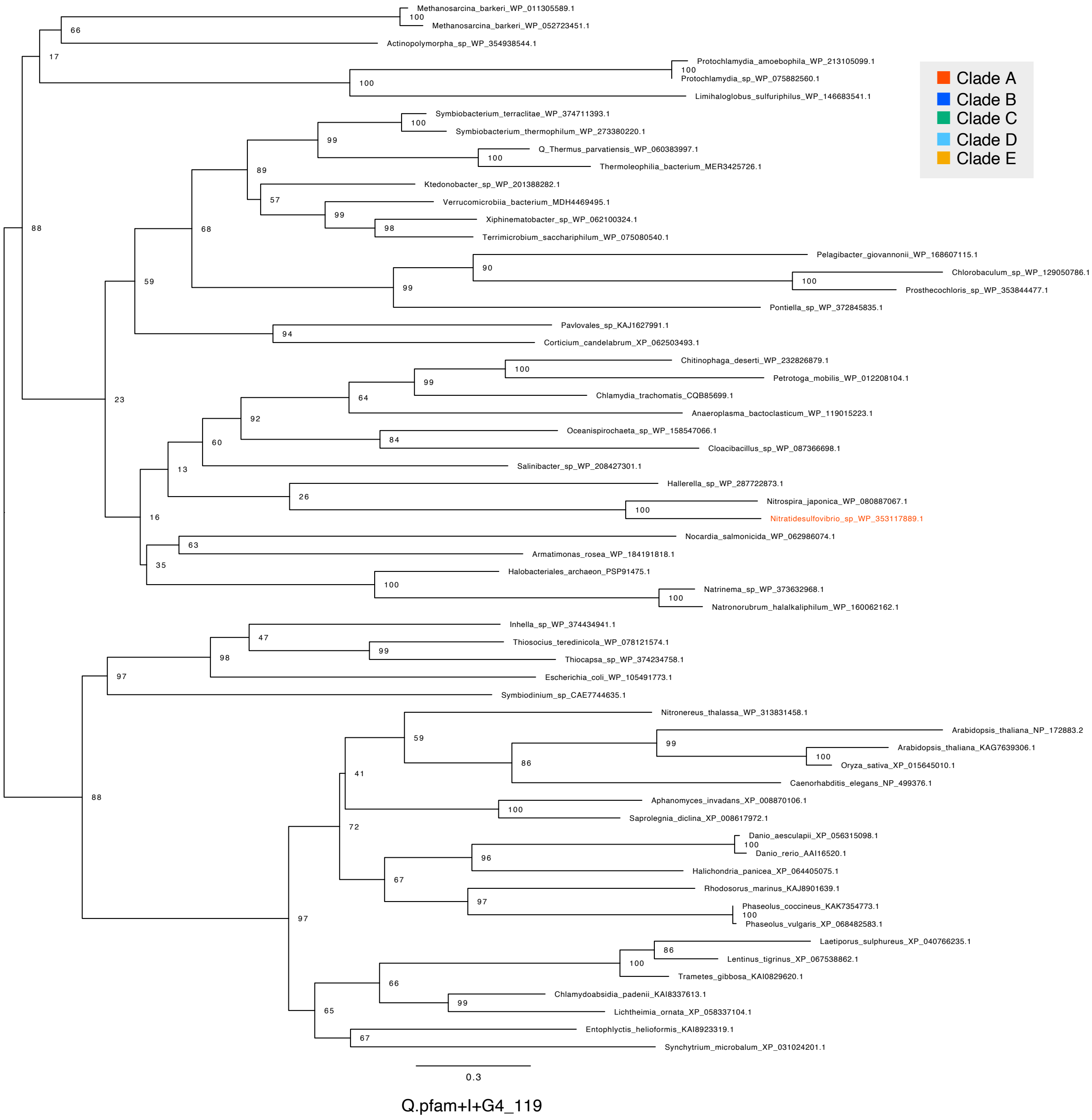

【Fig. S6C. Maximum likelihood phylogenetic tree of iPSP1.】

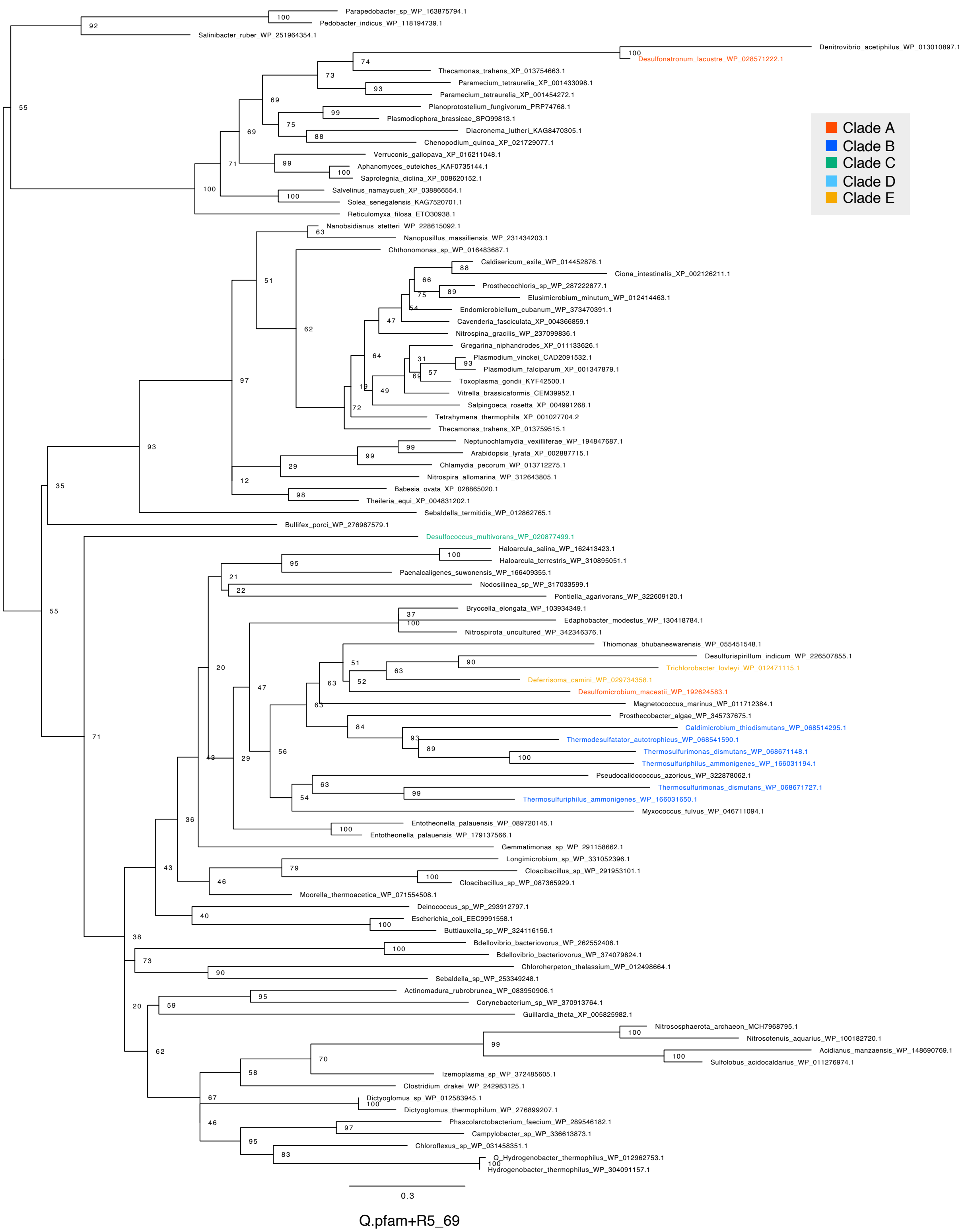


**Supplementary Figure 6. Maximum likelihood phylogenetic tree of PSP**

(A) dPSP1; (B) dPSP2; (C) iPSP1. Bootstrap values were calculated using the ultrafast bootstrap method. The evolutionary model, scale bar and number of sites are shown in the figure. Sequence in the phylum TDB are highlighted with colors.

【Fig. S7. Maximum likelihood phylogenetic tree of SHMT.】

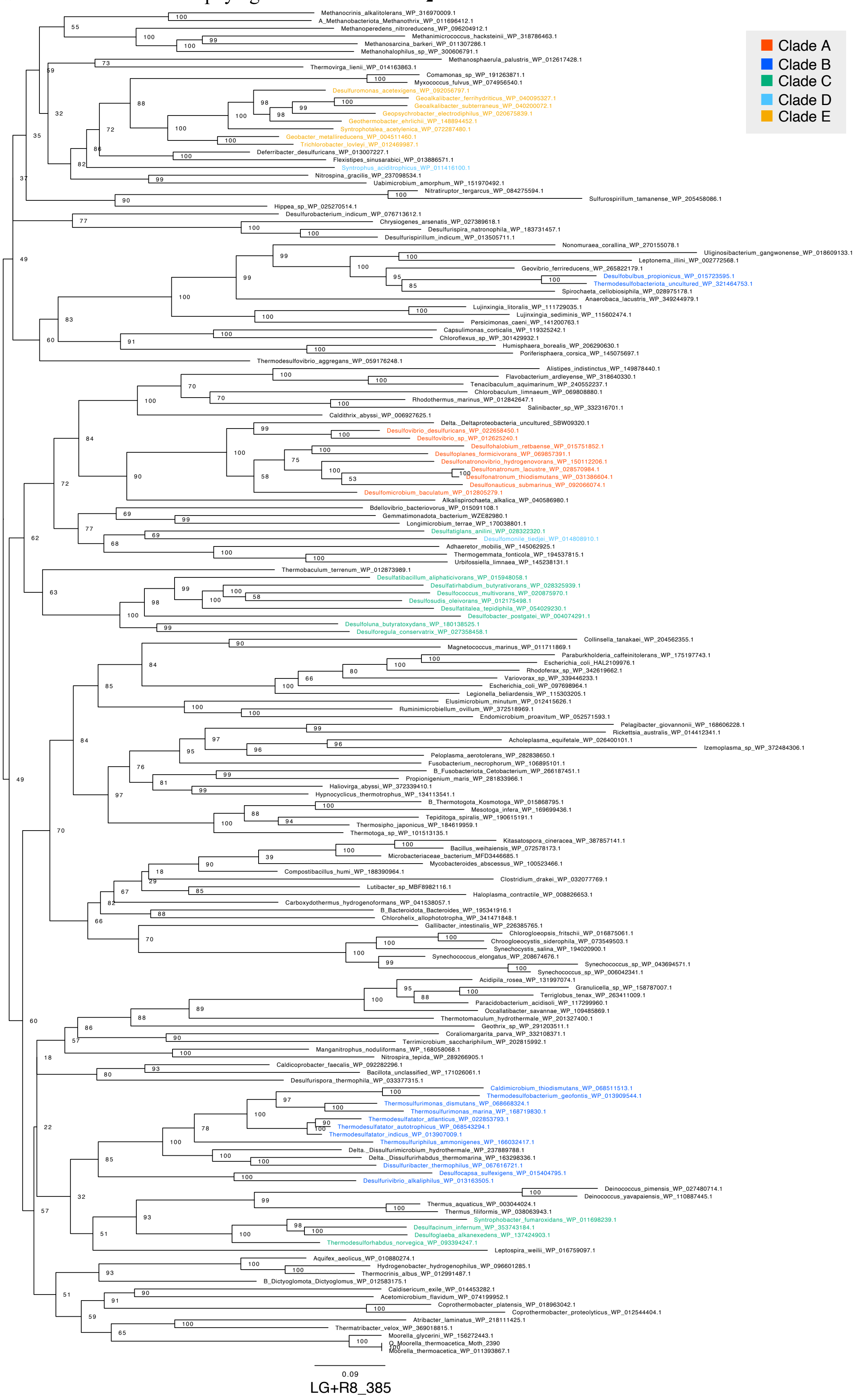

**Supplementary Figure 7. Maximum likelihood phylogenetic tree of SHMT.**
Bootstrap values were calculated using the ultrafast bootstrap method. The evolutionary model, scale bar and number of sites are shown in the figure. Sequence in the phylum TDB are highlighted with colors.

【 Fig. S8A. Maximum likelihood phylogenetic tree of GCST.】

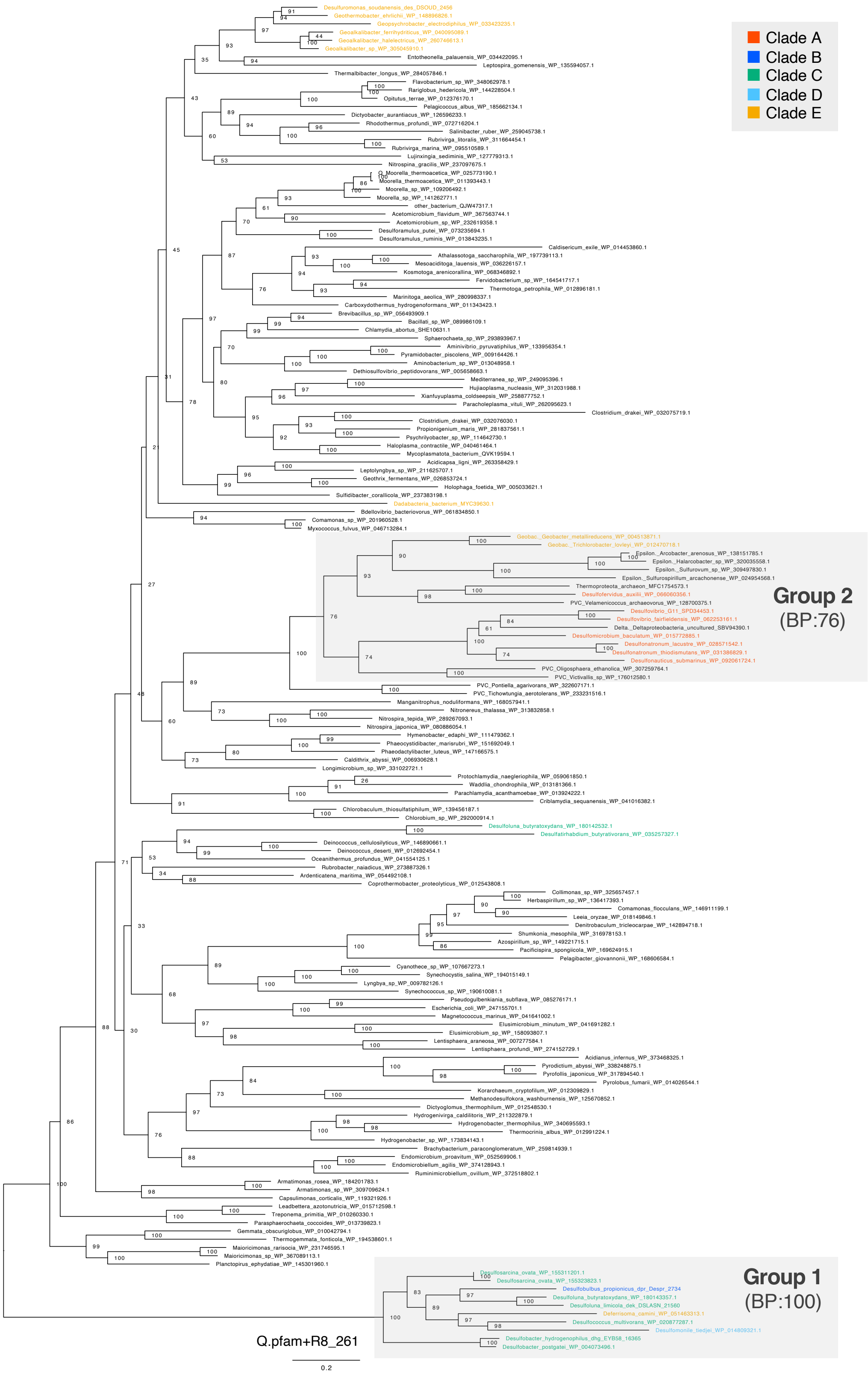

【Fig. S8B. Maximum likelihood phylogenetic tree of GCSL.】

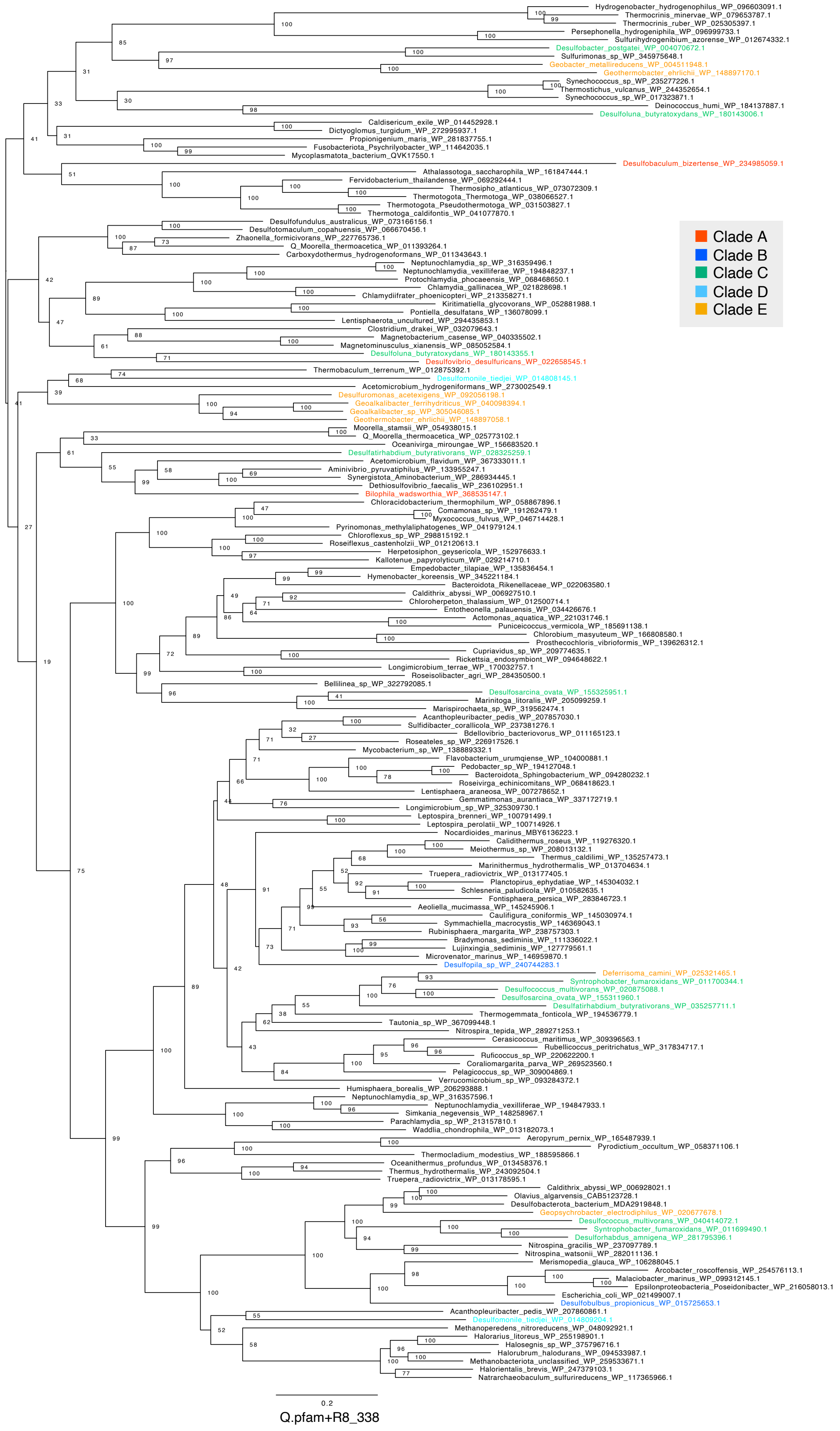

【 Fig. S8C. Maximum likelihood phylogenetic tree of GCSH.】

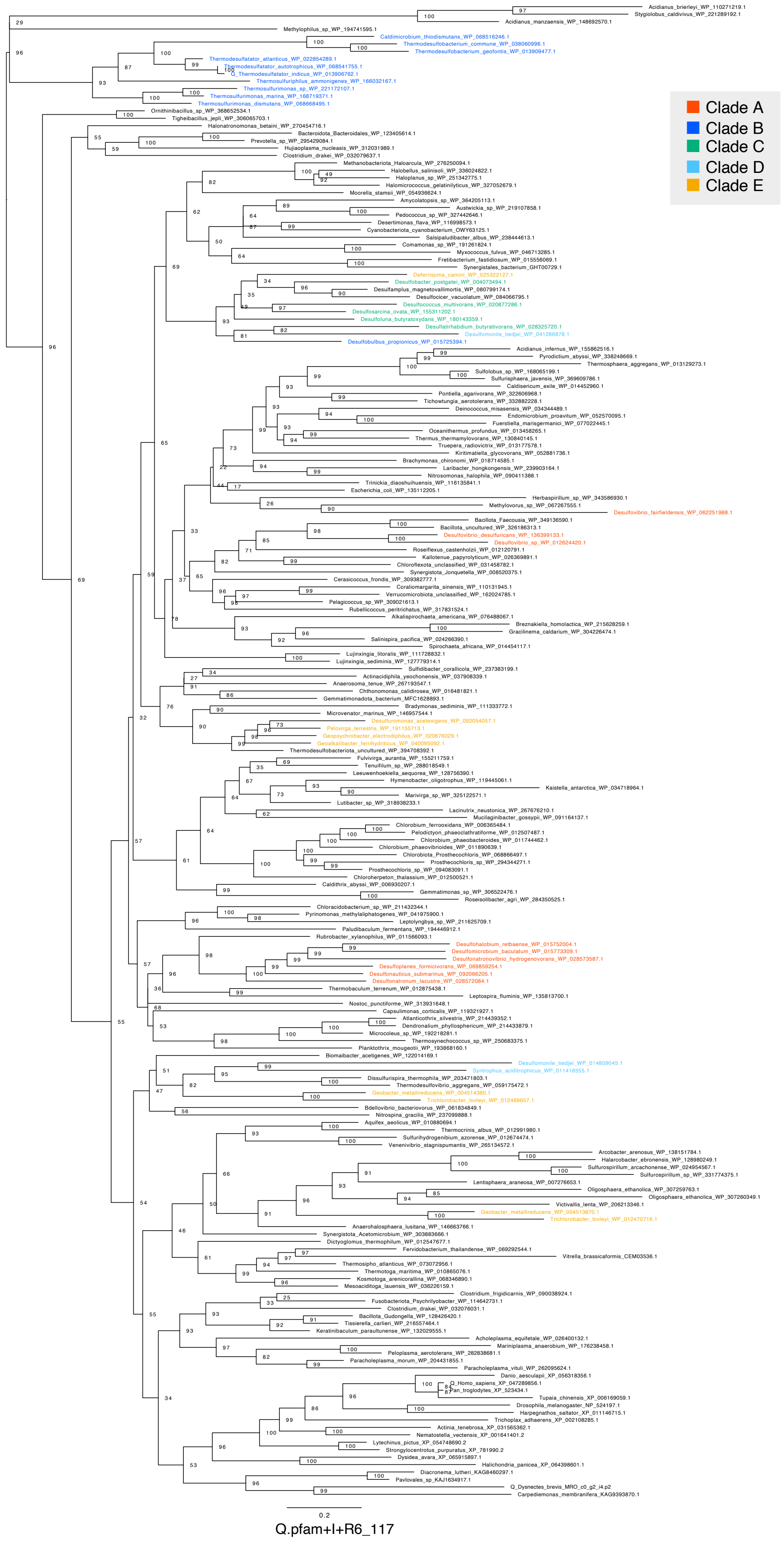

**Supplementary Figure 8. Maximum likelihood phylogenetic tree of the GCS complex.**
(A) GCST; (B) GCSL; (C) GCSH. Bootstrap values were calculated using the ultrafast bootstrap method. The evolutionary model, scale bar and number of sites are shown in the figure. Sequence in the phylum TDB are highlighted with colors.

【Fig. S9A. Maximum likelihood phylogenetic tree of GR-A.】

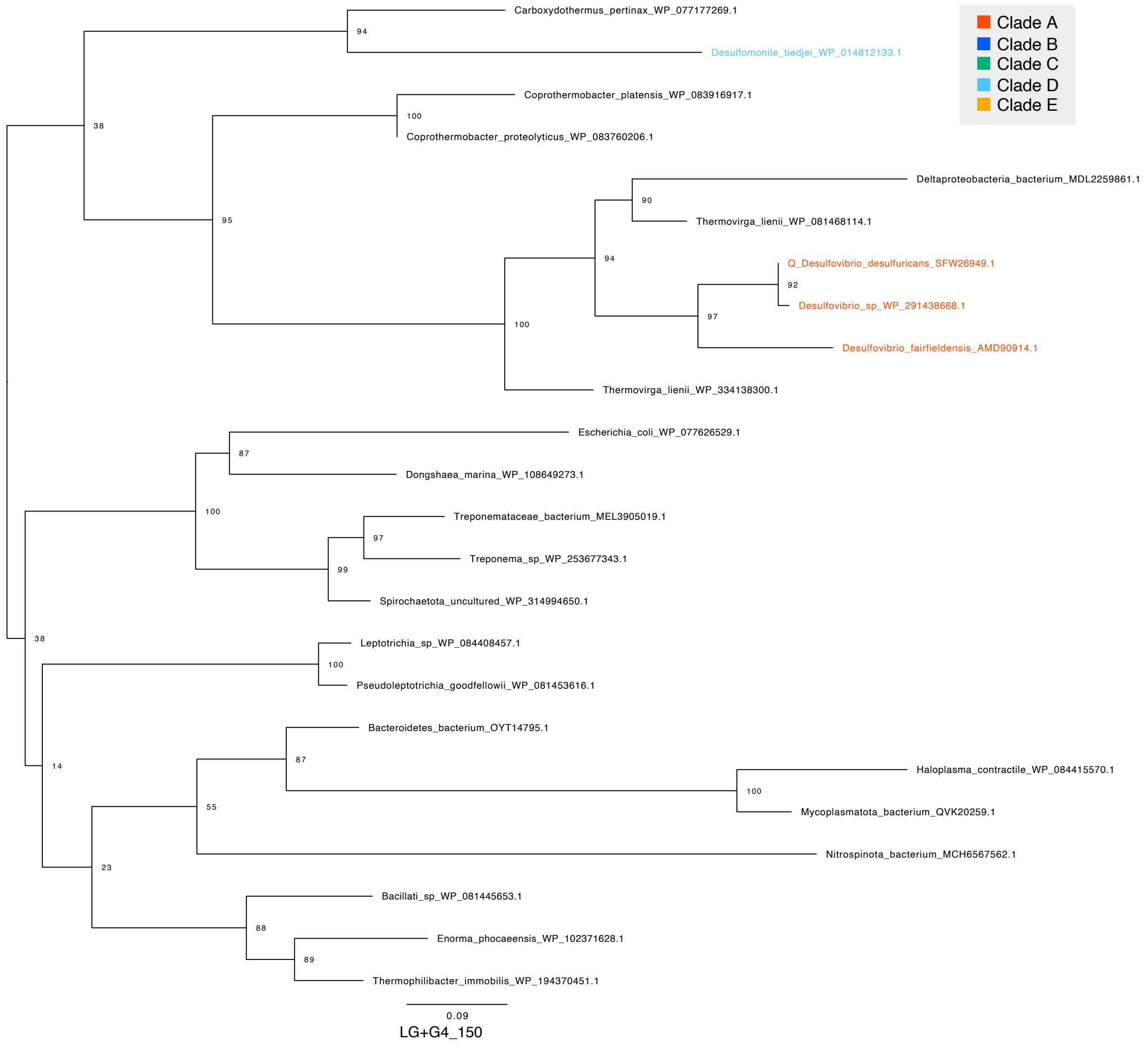

【 Fig. S9B. Maximum likelihood phylogenetic tree of GR-Bab.】

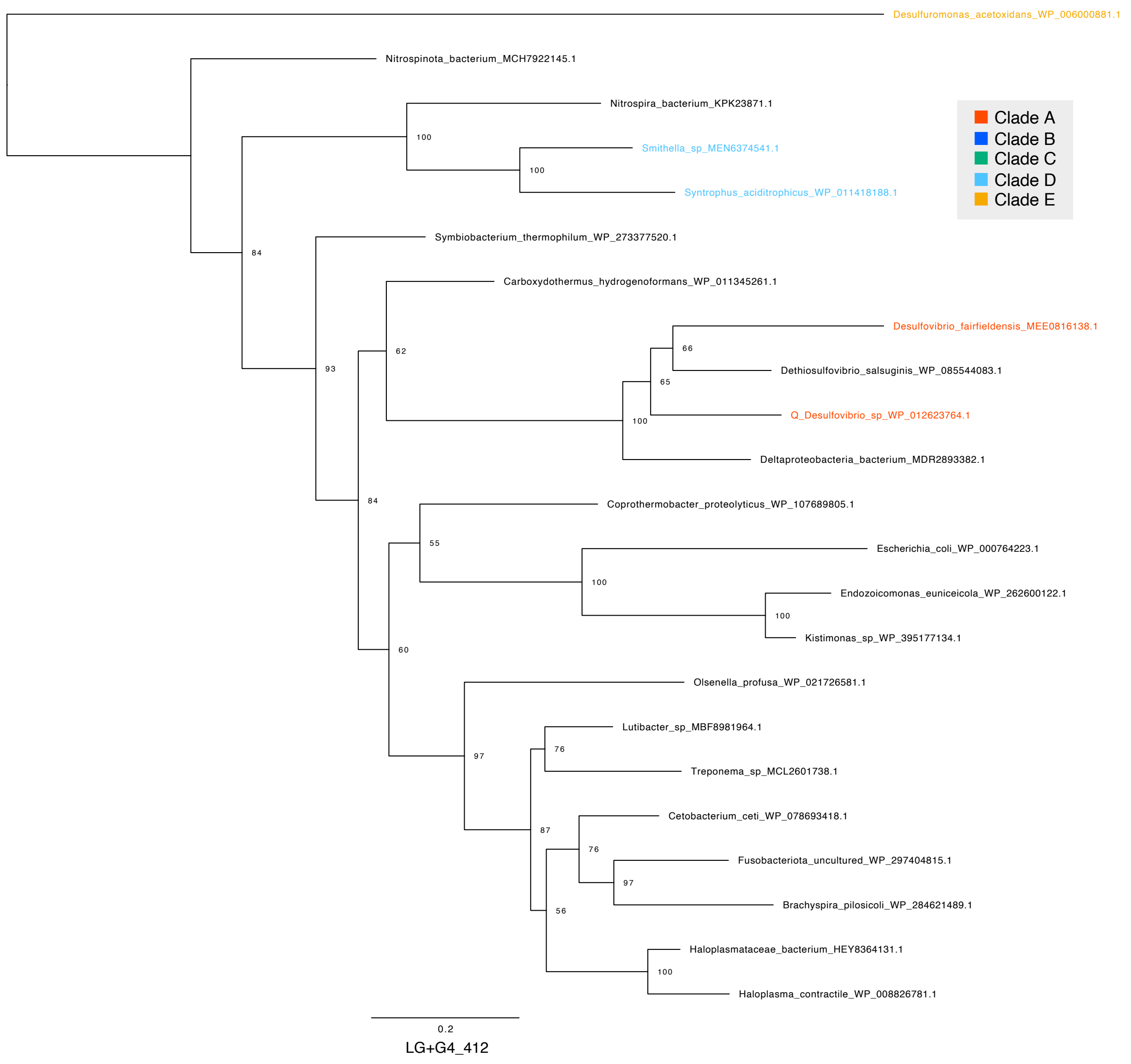

【 Fig. S9C. Maximum likelihood phylogenetic tree of GR-Bg.】

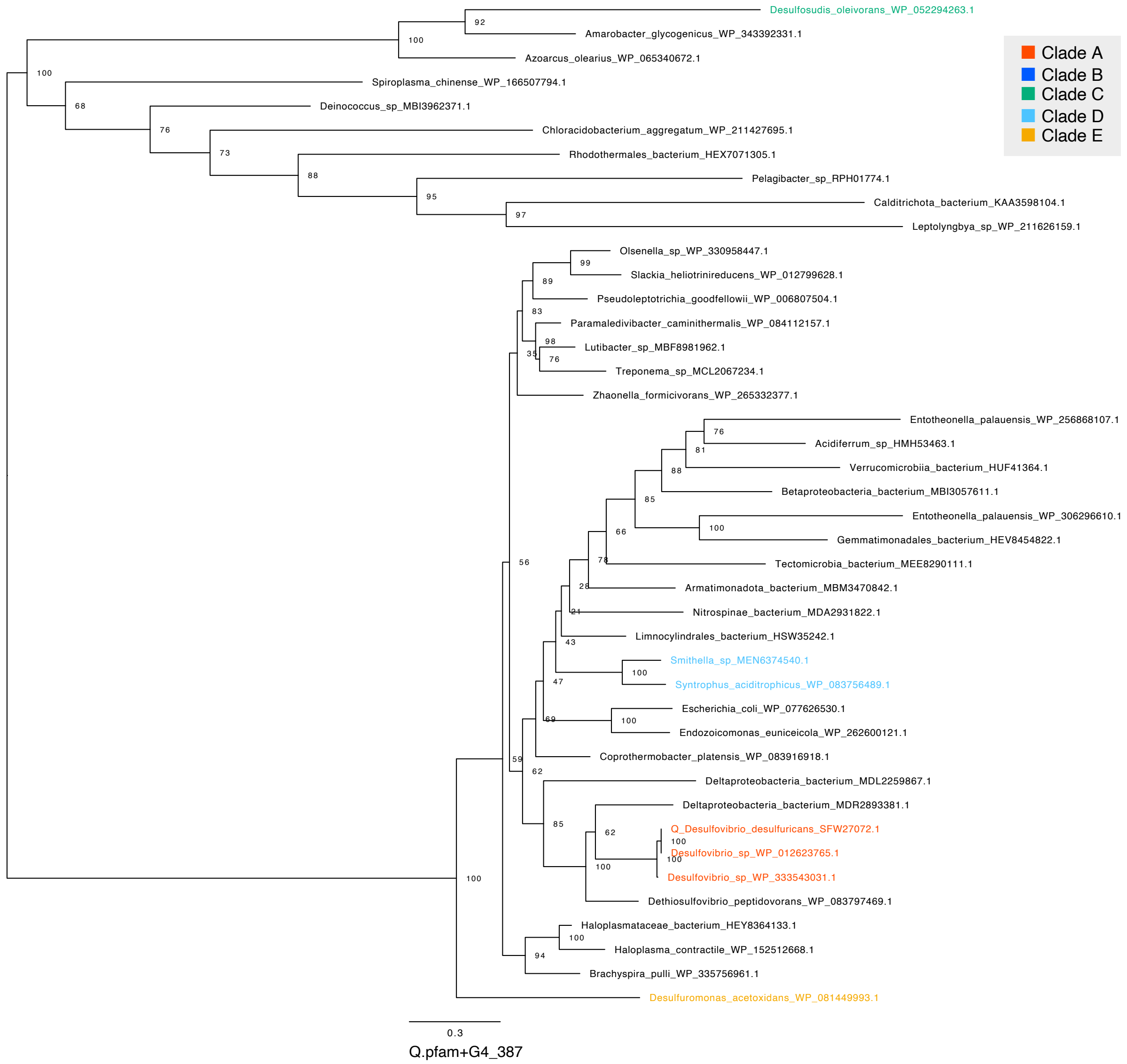

【 Fig. S9D. Maximum likelihood phylogenetic tree of GR-Ca.】

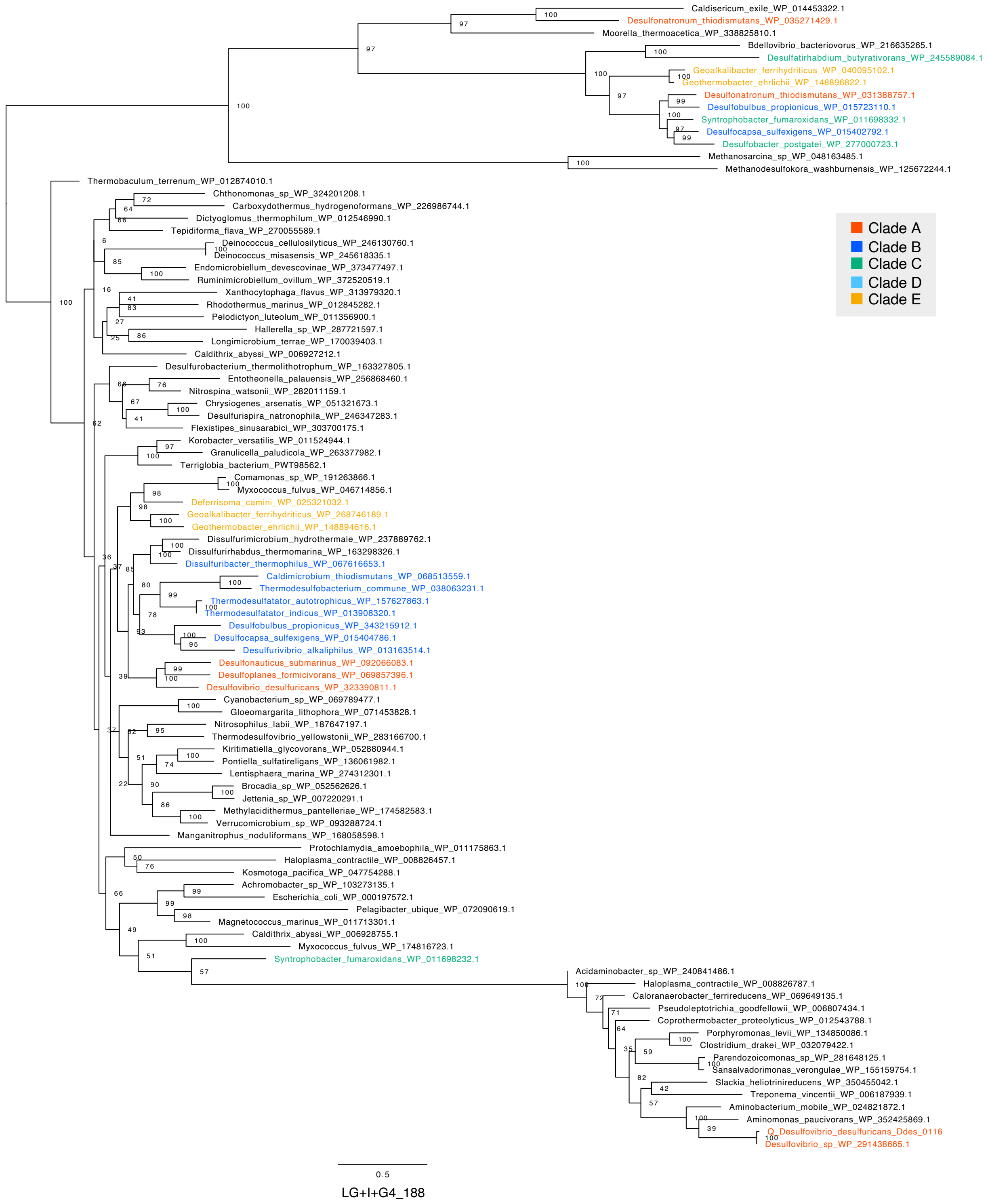

【 Fig. S9E. Maximum likelihood phylogenetic tree of GR-Cb.】

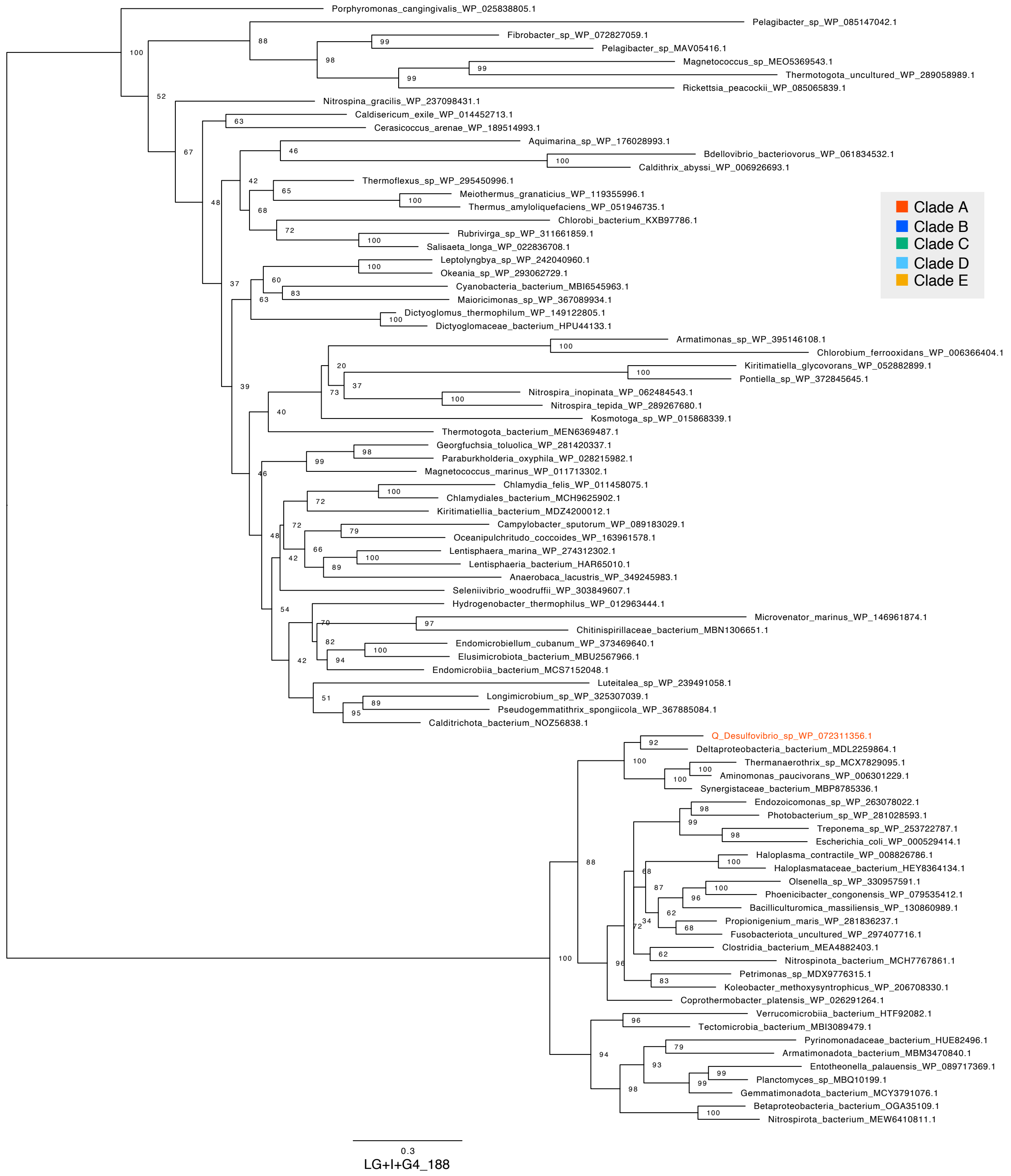


**Supplementary Figure 9. Maximum likelihood phylogenetic tree of the GR.**
(A) GR-A; (B) GR-Bab; (C) GR-Bg; (D) GR-Ca; (E) GR-Cb. Bootstrap values were calculated using the ultrafast bootstrap method. The evolutionary model, scale bar and number of sites are shown in the figure. Sequence in the phylum TDB are highlighted with colors.

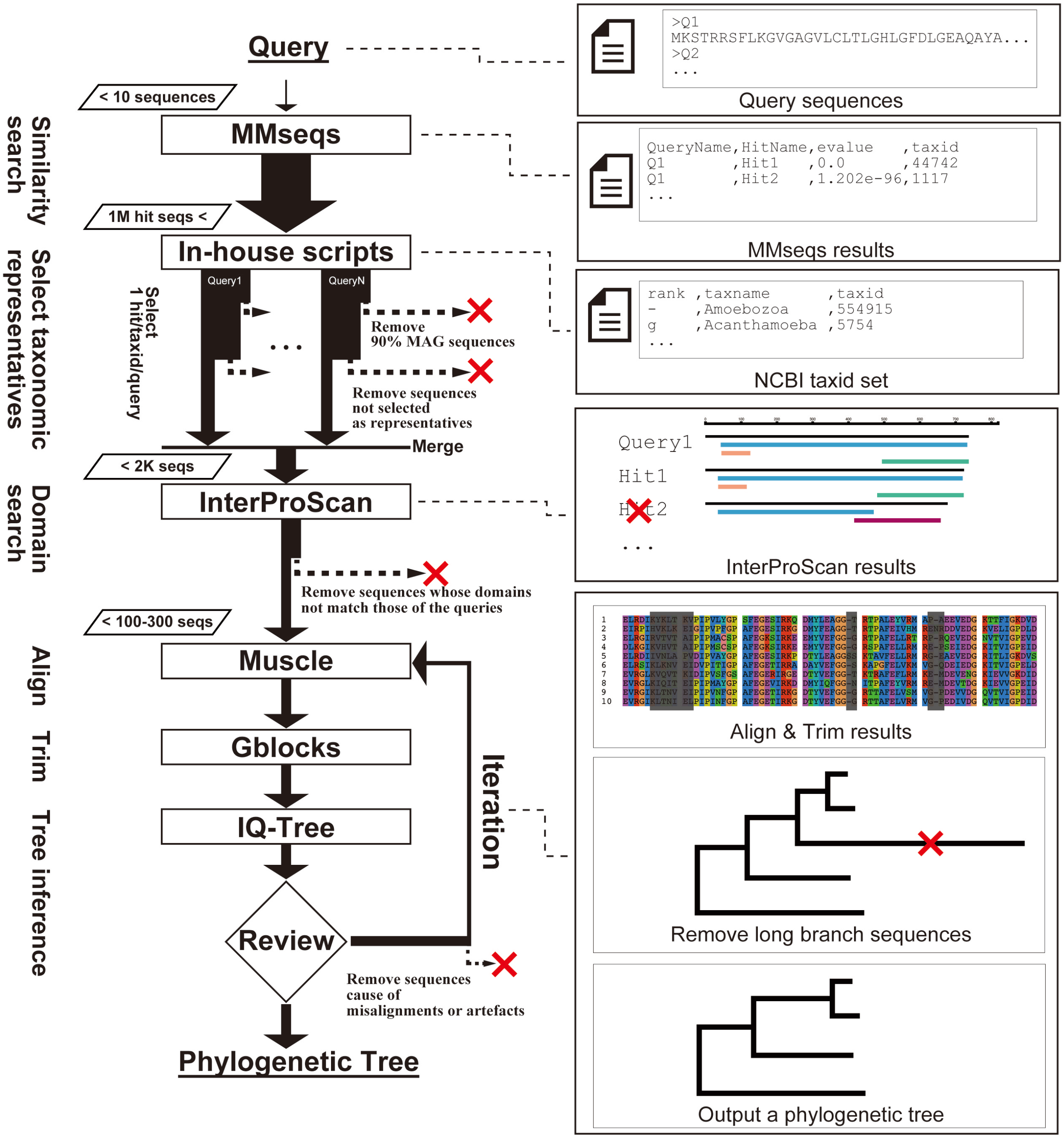


**Supplementary Figure 8. Workflows for phylogenetic analysis.**
